# The Arrival of Steppe and Iranian Related Ancestry in the Islands of the Western Mediterranean

**DOI:** 10.1101/584714

**Authors:** Daniel M. Fernandes, Alissa Mittnik, Iñigo Olalde, Iosif Lazaridis, Olivia Cheronet, Nadin Rohland, Swapan Mallick, Rebecca Bernardos, Nasreen Broomandkhoshbacht, Jens Carlsson, Brendan J. Culleton, Matthew Ferry, Beatriz Gamarra, Martina Lari, Matthew Mah, Megan Michel, Alessandra Modi, Mario Novak, Jonas Oppenheimer, Kendra A. Sirak, Kirstin Stewardson, Stefania Vai, Edgard Camarós, Carla Calò, Giulio Catalano, Marian Cueto, Vincenza Forgia, Marina Lozano, Elisabetta Marini, Margherita Micheletti, Roberto M. Miccichè, Maria R. Palombo, Damià Ramis, Vittoria Schimmenti, Pau Sureda, Luís Teira, Maria Teschler-Nicola, Douglas J. Kennett, Carles Lalueza-Fox, Nick Patterson, Luca Sineo, David Caramelli, Ron Pinhasi, David Reich

## Abstract

A series of studies have documented how Steppe pastoralist-related ancestry reached central Europe by at least 2500 BCE, while Iranian farmer-related ancestry was present in Aegean Europe by at least 1900 BCE. However, the spread of these ancestries into the western Mediterranean where they have contributed to many populations living today remains poorly understood. We generated genome-wide ancient DNA from the Balearic Islands, Sicily, and Sardinia, increasing the number of individuals with reported data from these islands from 3 to 52. We obtained data from the oldest skeleton excavated from the Balearic islands (dating to ∼2400 BCE), and show that this individual had substantial Steppe pastoralist-derived ancestry; however, later Balearic individuals had less Steppe heritage reflecting geographic heterogeneity or immigration from groups with more European first farmer-related ancestry. In Sicily, Steppe pastoralist ancestry arrived by ∼2200 BCE and likely came at least in part from Spain as it was associated with Iberian-specific Y chromosomes. In Sicily, Iranian-related ancestry also arrived by the Middle Bronze Age, thus revealing that this ancestry type, which was ubiquitous in the Aegean by this time, also spread further west prior to the classical period of Greek expansion. In Sardinia, we find no evidence of either eastern ancestry type in the Nuragic Bronze Age, but show that Iranian-related ancestry arrived by at least ∼300 BCE and Steppe ancestry arrived by ∼300 CE, joined at that time or later by North African ancestry. These results falsify the view that the people of Sardinia are isolated descendants of Europe’s first farmers. Instead, our results show that the island’s admixture history since the Bronze Age is as complex as that in many other parts of Europe.

## Introduction

The advent of the European Bronze Age after 3000 BCE was marked by an increase in long-range human mobility. People with ancestry from the Steppe north of the Black and Caspian Seas made a profound demographic impact in central and eastern Europe, mixing with local farmers to contribute up to three quarters of the ancestry of peoples associated with the Corded Ware complex^1–3^. The expansion of the Beaker complex after around 2400 BCE from the west had a less straightforward correlation to genetic ancestry. In Iberia, most people buried with artifacts of the Beaker complex had little if any Steppe pastoralist-related ancestry (from here on denoted “Steppe ancestry”), but Beaker cultural practices were adopted by people in Central Europe were in part descended from Steppe pastoralists and then spread this material culture along with Steppe ancestry to northwestern Europe^4^. In Iberia, Steppe ancestry began to appear in outlier individuals by ∼2500 BCE^4^, and became fully mixed into the Iberian population by 2000 BCE^5^. Meanwhile on Crete in the eastern Mediterranean, there was little if any Steppe ancestry identified in all published samples from the Middle to Late Bronze Age “Minoan” culture (individuals dating to 2400-1700 BCE), although these individuals derived about 15% of their ancestry from groups related to early Iranian farmers (from here on referred to as “Iranian-related ancestry”)^6^ (**Fig. 1**).

**Figure 1:**
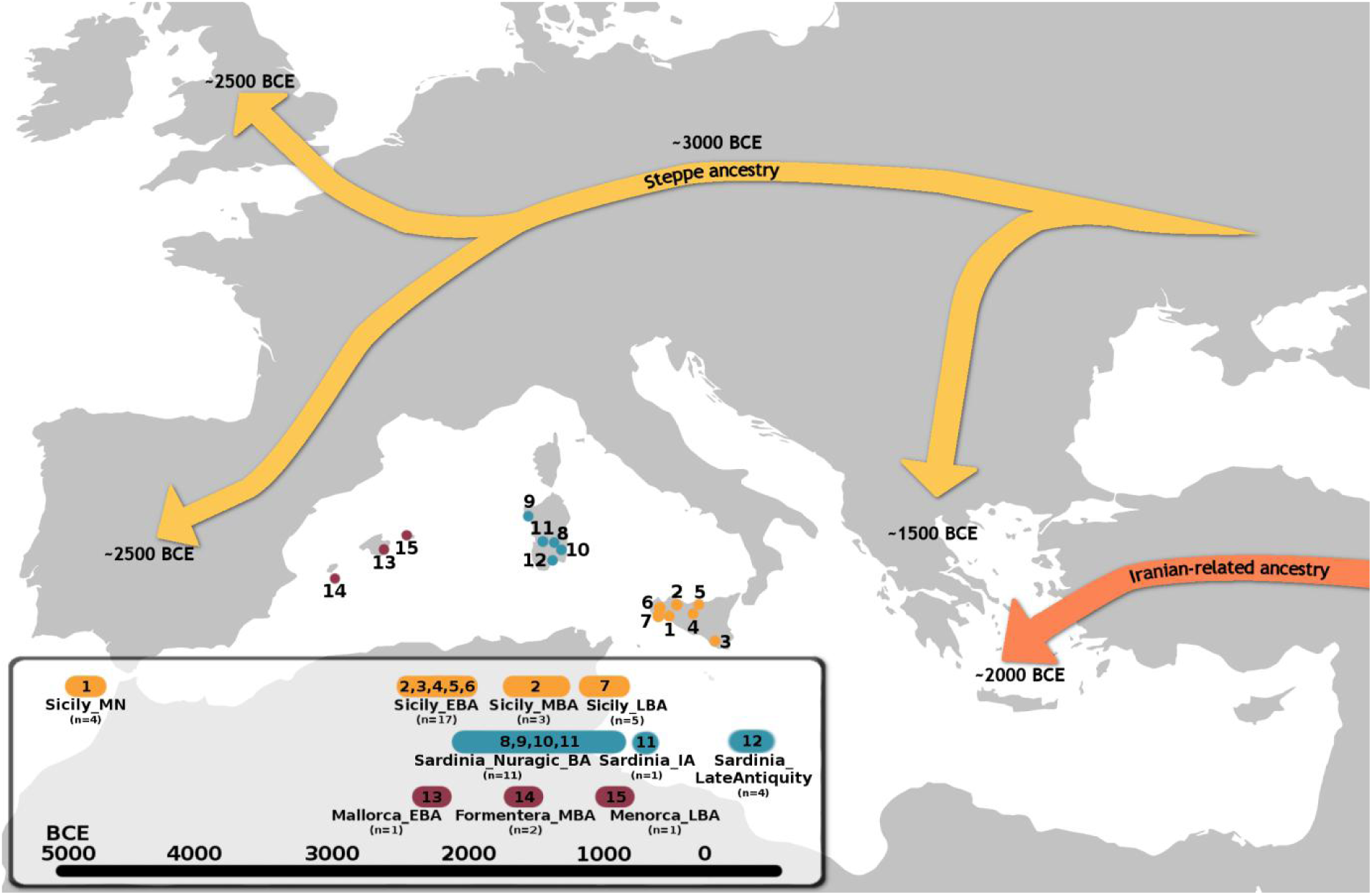
Timeline and geographical origins of the 49 newly reported ancient individuals along with the previously reported individual for whom we increase data quality. 1-Fossato di Stretto Partana; 2-Buffa cave; 3-Contrada Paolina; 4-Isnello; 5-Vallone Inferno; 6-Marcita; 7-Salaparuta; 8-Seulo; 9-Alghero-Lu Maccioni cave; 10-Perdasdefogu; 11-Usellus; 12-Grotta Colombi; 13-Cova des Moro; 14-Cap de Barbaria; 15-Naveta des Tudons.

In the islands of the central and western Mediterranean, the Bronze Age transition has not been investigated with ancient DNA, despite the fact that archaeological evidence reveals that many of the same cultural changes that affected mainland Europe and the eastern Mediterranean also impacted this region^7^. The first evidence for a permanent human presence in the Balearic Islands is dated to just before the onset of the Bronze Age in this part of Europe, between ∼2500-2300 BCE^8,9^. Early settlers initially relied on animal husbandry and their economy was focused on sheep, goat^9,10^, and cereal agriculture^11^, while exploitation of wild marine resources (fish, marine birds, mollusks) was central to subsistence on the small island of Formentera^10,12^. Around 1200 BCE, the development of the Talaiotic culture in Mallorca and Menorca (the easternmost Balearic Islands) was marked by intensified management of food resources and the appearance of monumental towers, the eponymous talaiots. These structures were similar in style to the Sardinian nuraghi^10,13^, raising the question of whether there was a cultural connection^14^, a scenario that would gain plausibility if there was substantial genetic exchange between the two regions. Nuragic Sardinians were also in cultural contact with groups from the eastern Mediterranean^15^, so an important question is whether they were admixed with either Steppe or Iranian-related ancestry. Meanwhile, the central Mediterranean island of Sicily was affected by the spread of Beaker cultural complex after around 2400 BCE, and by cultural influence from the Aegean in the Late Helladic Period ∼1600-1200 BCE (the period of the “Mycenaean” culture)^16–18^. An unanswered question is whether these events or other cultural changes on the island involved substantial movements of people.

We increased the number of individuals from these islands with genome-wide data from 3 to 52, and analyzed the data to address three questions. First, to what extent did movements of people into these islands track the material culture exchanges documented in the archaeological record? Secondly, can we establish the source and minimum dates of arrival of Steppe ancestry in the central and western Mediterranean islands where this ancestry is present in variable proportions today? Thirdly, did Iranian-related ancestry reach the central and western Mediterranean prior to the period of Phoenician and Greek expansion?

## Results

### Samples and sequencing results

We prepared powder from petrous bones and teeth in dedicated ancient DNA clean rooms at University College Dublin, Harvard Medical School, and the University of Florence, extracted DNA using a method designed to retain short molecules^19–22^, and converted the extracted DNA into double-stranded libraries^23^. We treated all libraries with Uracil-DNA Glycosylase (UDG) to cleave the analyzed molecules at damaged uracil sites, thereby greatly reducing the rate of cytosine-to-thymine errors characteristic of ancient DNA. We enriched ancient DNA libraries for sequences overlapping approximately 1.24 million single nucleotide polymorphisms (SNPs)^24,25^, and obtained genome-wide data from a total of 49 individuals from the Balearic Islands, Sardinia, and Sicily while increasing the quality of data for a Bell Beaker culture associated individual from Sicily (adding three more libraries to the one previously generated) (**Fig. 1, Online Table 1, Supplementary Materials**). We established chronology based on archaeological context and by assembling direct radiocarbon dates on bone for 28 of the individuals (direct dates for 26 individuals are reported for the first time here; **Online Table 2**). We removed from the analysis dataset eight individuals with fewer than 20,000 of the targeted SNPs covered by at least one sequence, and five with evidence of substantial contamination or less than 3% cytosine-to-thymine error in terminal cytosines. We also removed one individual who we detected as a first degree relative (a son) of another (his mother) that gave higher quality genetic data. This left 36 individuals for our modeling (however, since all the data are useful we fully report all individuals, **Online Table 1**). In the analysis dataset, the median coverage on targeted SNPs on chromosomes 1-22 was 3.20-fold (range 0.02-12.13), and the median number of SNPs covered by at least one sequence was 756709 (range 23600-1038409). All mitochondrial DNA point estimates for match rate to the consensus sequence had 95% confidence intervals with upper bounds from 0.96-1.00, while contamination estimates based on X chromosome variation (meaningful only in males) were all below 1.1% (**Online Table 1**). All individuals had data from at least one library with cytosine-to-thymine damage in the terminal nucleotides greater than 3% (the minimum suggested as a guideline for the plausible authentic DNA^23^). The qualitative patterns of ancestry in the data were unchanged when we restricted to transversion SNPs which are not affected by characteristic ancient DNA errors (**Supplementary Fig. 1**).

### Genetic affinities and population groupings

We carried out principal component analysis (PCA) of the ancient individuals merged with previously published ancient DNA^1,2,4,6,26–44^, projected onto genetic variation among 737 diverse present-day west Eurasians genotyped at ∼600,000 SNPs (a subset of the positions on the ∼1.24 million SNP set)^27,39,45–47^ (**Fig. 2b, Online Table 3**). We also performed unsupervised clustering with ADMIXTURE^48^ (**Fig. 2a**). The three Balearic Islands individuals in the analysis dataset fall between the European Neolithic and Bronze Age clusters on the PCA, consistent with harboring Steppe ancestry (**Fig. 2b**), a finding that is also supported by the finding in these individuals of an ADMIXTURE component maximized in Eastern European Hunter Gatherers (*EHG*) and Yamnaya Steppe Pastoralists. The eight Nuragic Sardinians cluster in PCA and ADMIXTURE with Middle Neolithic Europeans, with the exception of one individual (I10365: 1643-1263 calBCE) that shows a shift towards the Sicilian cluster. The Iron Age Sardinian (∼400-200 BCE) and the four Late Antiquity Sardinians (∼200-700 CE) deviate toward the Mycenaean cluster, while one of the Late Antiquity Sardinians also deviates toward Central European Bronze Age individuals. The 20 Sicilians cluster in PCA and ADMIXTURE mostly with the European Neolithic individuals, with the exception of two that have more affinity to the Central European Bronze Age individuals (**Fig. 2**). Relative to the Middle Neolithic Sicilians (*Sicily_MN*), the main Bronze Age Sicilian cluster (after removing these outliers) deviates in a more subtle way toward eastern groups (either Steppe pastoralists or individuals from the Aegean Bronze Age), a pattern that is also evident in ADMIXTURE.

**Figure 2:**
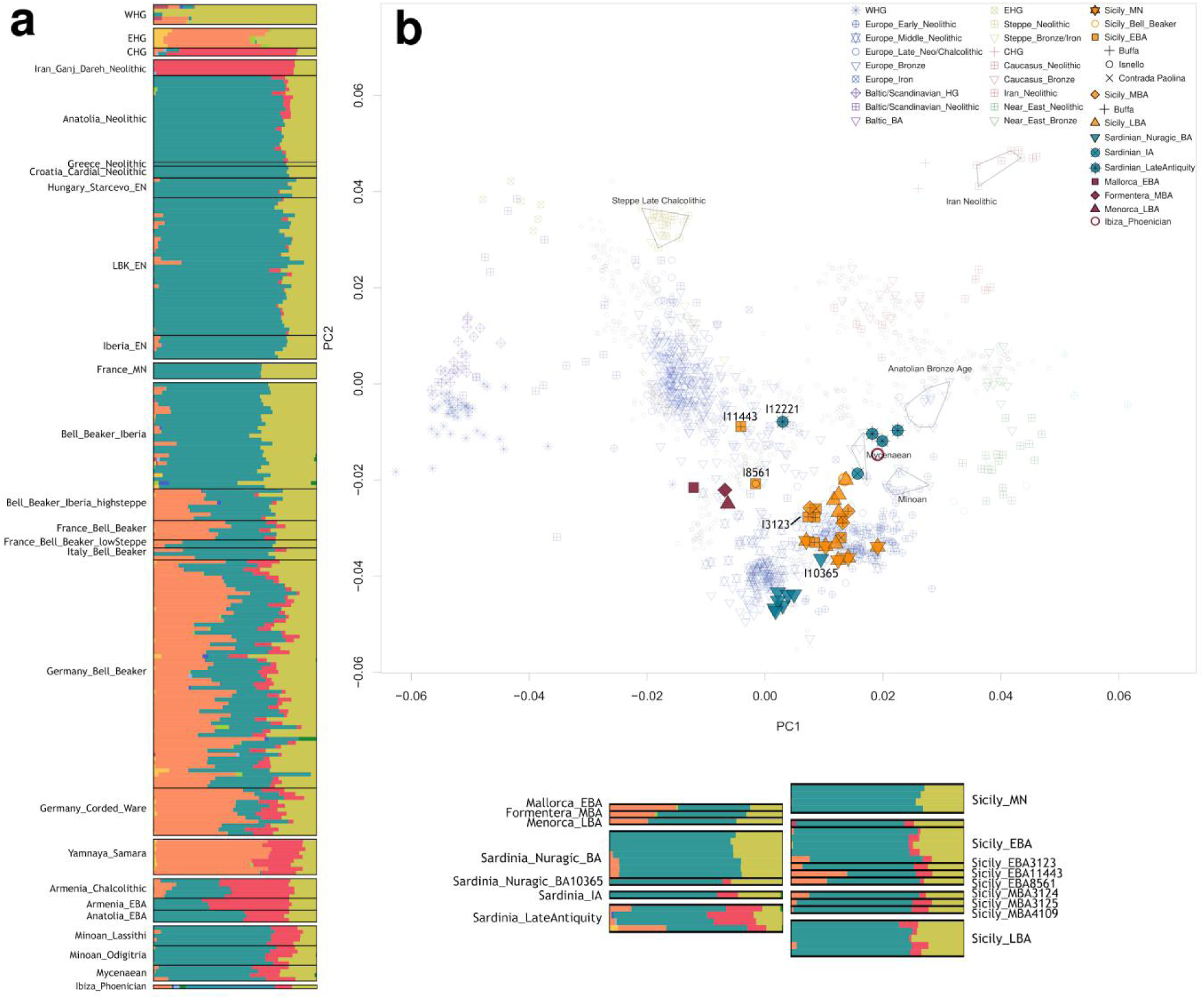
Ancestry of ancient Sardinians, Sicilians and Balearic islanders and other ancient and present-day populations according to a) unsupervised ADMIXTURE analysis with K=10 clusters; and b) PCA with previously published ancient individuals (non-filled symbols), projected onto variation from present-day populations (gray squares).

To formally cluster these individuals, we used *qpWave*^*2*^ to test whether each individual in turn was consistent with being from the same group as others from the same time period and region (that is, we tested whether they were consistent with forming a clade at a p<0.010 level) (**Supplementary Materials, Fig. 3**). In some instances where the *qpWave* results were ambiguous, we carried out more refined tests to split individuals into analysis groupings **(Supplementary Materials).**

**Figure 3:**
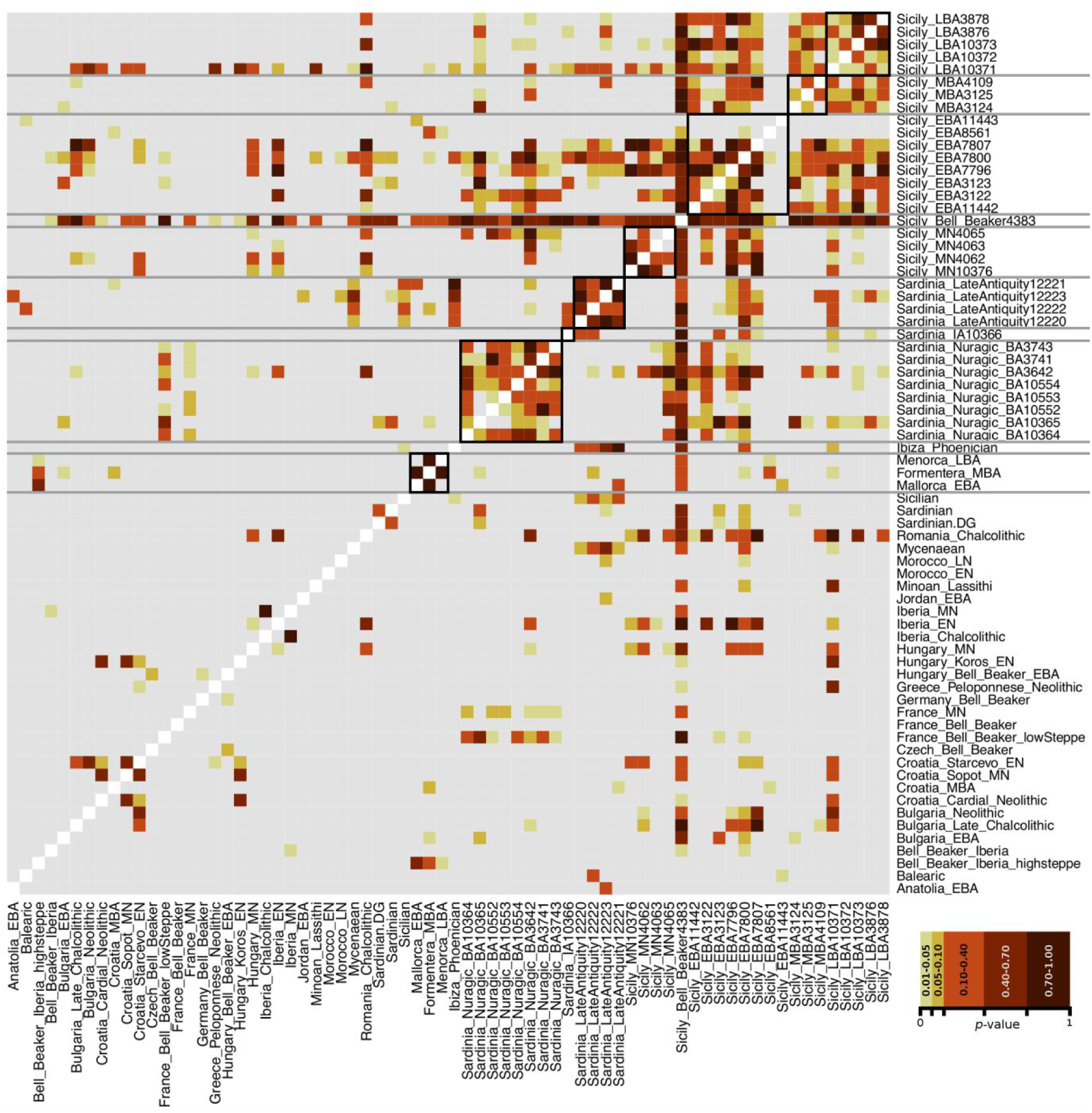
Pairwise *qpWave* testing to group individuals. Black lines represent the initial clusters of individuals from this study by location and/or period. Gray-coloured models have a p-value below 0.010 and are rejected.

In the Balearic Islands, *qpWave* revealed significant differences between the Early Bronze Age individual *Mallorca_EBA* and the Late Bronze Age individual *Menorca_LBA* (p=0.002) (**Supplementary Table 1**). While *qpWave* tests comparing the Middle Bronze Age individual *Formentera_MBA* to the other two individuals were non-significant, the symmetry test statistic *f*_*4*_(*Mbuti.DG, Iberia_Chalcolithic*; *Formentera_MBA, Menorca_LBA*) was Z = 2.6 standard errors from zero (exceeding a threshold of |Z|>2, which is approximately p<0.05), implying significantly different ancestry in *Formentera_MBA* than in *Menorca_LBA*. In light of this and the different dates and island sources of these three individuals, we treated all three separately for analysis.

For the Nuragic Sardinians (*Sardinia_Nuragic_BA*), only I10365 clearly did not form a clade with others of the same cultural affiliation (**Supplementary Tables 2 and 3**). Thus, we treated this individual whose radiocarbon date confirms it as contemporaneous with the others as an outlier (*Sardinia_Nuragic_BA10365*). The Iron Age Sardinian individual formed a clade with some from the *Sardinian_LateAntiquity* cluster and some ancient Sicilians, but we treated it separately because of its distinctive time period and geographic location. The four *Sardinian_LateAntiquity* individuals were consistent with forming a clade in *qpWave*, but one individual separated from the others in PCA (**Fig. 2**), and also showed distinct signals in admixture modeling, and hence we analyzed it separately as *Sardinian_LateAntiquity12221* (**Supplementary Table 4**).

For Sicily, our analysis confirmed the two Early Bronze Age outliers *Sicily_EBA11443* and *Sicily_EBA8561* evident in PCA and ADMIXTURE (both p<10^-12^ relative to the main cluster), while identifying a third outlier *Sicily_EBA3123* (p=0.004) (**Supplementary Tables 5 and 6**). One Sicilian Middle Bronze Age individual was not consistent with being a clade with one of the other two, and we treated the three separately in subsequent analysis (*Sicily_MBA3124, Sicily_MBA3125*, and *Sicily_MBA4109*) (**Supplementary Table 7**). All 5 Late Bronze Age individuals were consistent with being a clade at the p>0.01 threshold and we grouped them (*Sicily_LBA*) (**Supplementary Table 8**).

We used *qpAdm*^*2,45*^ to decompose the ancestry of each analysis grouping into four “distal” sources: *Anatolia_Neolithic*, Western Hunter-Gatherers (*WHG*), *Iran_Ganj_Dareh_Neolithic* and *Yamnaya_Samara*. We first tested the model with *Anatolia_Neolithic* and *WHG*, then added either *Iran_Ganj_Dareh_Neolithic* or *Yamnaya_Samara* as a potential third source, and finally combined all ancestry sources for a total of four sources. We quote the most parsimonious model (as measured by the lowest number of ancestry sources) that fits at p>0.05. A unique parsimonious model fit for each analysis grouping (**Fig. 4b and Supplementary Table 9 and 10**).

**Figure 4:**
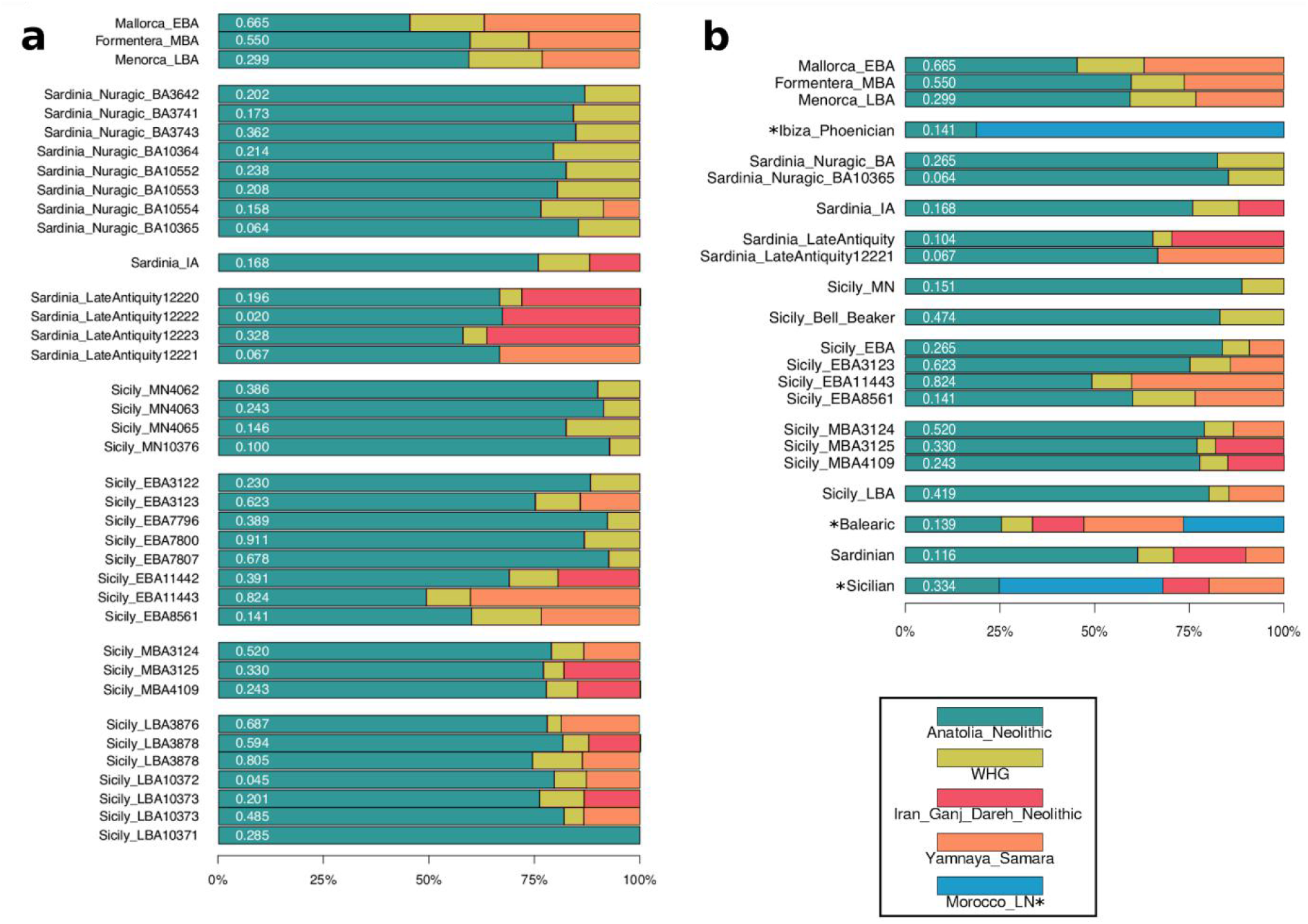
Proportions of ancestry using a distal *qpAdm* framework on an individual basis (a), and based on *qpWave* clusters (Fig. 3) (b). We show all valid models (p>0.05) for the lowest possible ranks. Some individuals produced two valid models at p>0.05, but for b), there is only a single parsimonious model for each analysis grouping. In panel b) relevant published individuals (*Ibiza_Phoenician, Sicily_Bell_Beaker*) and modern populations (*Balearic, Sardinian, Sicilian*) are also presented. Some of these did not produce valid models with the four base ancestries so we show the most parsimonious working models after including *Morocco_LN* (denoted by an asterisk). P-values in white are within bars (Supplementary Materials and Supplementary Tables 9 and 10 give all numbers underlying this figure).

### Formal modeling of the ancestry of Bronze Age Individual from the Balearic Islands

*Mallorca_EBA* dates to the earliest period of permanent occupation of the islands at around 2400 BCE^10,49^. We parsimoniously modeled *Mallorca_EBA* as deriving 36.9 ± 4.2% of her ancestry from a source related to *Yamnaya_Samara*; all fitting models require Steppe ancestry, whereas no Iranian-related ancestry is required to achieve a fit (**Fig. 4, Supplementary Table 9**). We next used *qpAdm* to identify “proximal” sources for *Mallorca_EBA*’s ancestry that are more closely related to this individual in space and time, and found that she can be modeled as a clade with the (small) subset of Iberian Bell Beaker culture associated individuals who carried Steppe-derived ancestry^4^ (p=0.442). This suggests that the movements of people that brought Steppe ancestry into Iberia may have been related to those that first settled the Balearic islands. However, archaeological evidence for the Beaker complex in the Balearic islands during the 3rd millennium BCE is scarce^9^, so it is possible that a related non-Beaker using group spread this ancestry.

Our estimates of Steppe ancestry in the two later Balearic Islands individuals are lower than the earlier one: 26.3 ± 5.1% for *Formentera_MBA* and 23.1 ± 3.6% for *Menorca_LBA* (**Supplementary Table 9**), but the Middle to Late Bronze Age Balearic individuals are not a clade relative to non-Balearic groups. Specifically, we find that *f*_*4*_(*Mbuti.DG*, X; *Formentera_MBA, Menorca_LBA*) is positive when X=*Iberia_Chalcolithic* (Z=2.6) or X=*Sardinia_Nuragic_BA* (Z=2.7). While it is tempting to interpret the latter statistic as suggesting a genetic link between peoples of the Talaiotic culture of the Balearic islands and the Nuragic culture of Sardinia, the attraction to *Iberia_Chalcolithic* is just as strong, and the mitochondrial haplogroup U5b1+16189+@16192 in *Menorca_LBA* is not observed in *Sardinia_Nuragic_BA* but is observed in multiple *Iberia_Chalcolithic* individuals. A possible explanation is that both the ancestors of Nuragic Sardinians and the ancestors of Talaiotic people from the Balearic Islands received gene flow from an unsampled Iberian Chalcolithic-related group (perhaps a mainland group affiliated to both) that did not contribute to *Formentera_MBA*.

During the Iron Age, Phoenician colonies were established in the Balearic islands. The Ibiza Phoenician individual published in ^50^ is not consistent with forming a clade with any of the Bronze Age individuals from the Balaeric islands newly reported in this study, and indeed we find that she can not be modeled even with our least parsimonious model of 4 distal sources. However, when we add in a North African source of ancestry, we can fit her as a two-way mix of 18.8 ± 7.9% *Anatolia_Neolithic* and 81.2 ± 7.9% *Morocco_LN* ancestry (p=0.141) (**Supplementary Materials**). We also can fit the Ibiza Phoenician as two-way mixture of a variety of groups closer to her in time one of which is always *Morocco_LN*. While several of these models include a Balaeric Island Bronze Age source, we cannot rule out the possibility that the Ibiza Phoenician individual has no local Balaeric ancestry at all. Specifically, we find that we can fit her with models that do not have a Balaeric source and that instead have Balaeric Bronze Age individuals in the outgroups (e.g. (e.g. 17.1 ± 3.5% *France_Bell_Beaker* and 82.9 ± 3.5% *Morocco_LN*, p=0.869) (**Supplementary Table 11**).

Modern Balearic individuals also do not fit with the least parsimonious model of 4 distal sources, however, we can fit them as a mixture of Steppe, Iranian-related, and North African ancestry, demonstrating the Balearic islands have been affected by significant admixture since the initial settlement.

### Formal Modeling of Ancestry Changes Over Time in Sardinia

We analyzed 13 individuals from Sardinia dated to ∼2200 BCE - 700 CE (**Fig. 1, Online Table 1**).

In *qpAdm*, all eight Bronze Age Nuragic individuals fit as descending from the same two deep ancestral sources (*Anatolia_Neolithic* and *WHG*), but mixed in different proportions: 82.5 ± 1.1% *Anatolia_Neolithic* for the main *Sardinia_Nuragic_BA* cluster (p=0.265), and 85.4 ± 2.2% for the *Sardinia_Nuragic_BA10365* outlier (p=0.064) (**Supplementary Table 9**). We find no working models when we consider chronological or geographically more proximal sources (e.g. Beaker complex associated individuals from Iberia, France, Czech Republic, Germany; or Chalcolithic Iberians and Neolithic Sicilians), although we do not have access to early Neolithic Sardinians for this analysis.

Most Sardinians buried in a Nuragic Bronze Age context possessed uniparental haplogroups found in European hunter-gatherers and early farmers, including Y-haplogroup R1b1a[xR1b1a1a] which is different from the characteristic R1b1a1a2a1a2 spread in association with the Bell Beaker complex^4^. An exception is individual I10553 (1226-1056 calBCE) who carried Y-haplogroup J2b2a (**Online Table 1**), previously observed in a Croatian Middle Bronze Age individual bearing Steppe ancestry^44^, suggesting the possibility of genetic input from groups that arrived from the east after the spread of first farmers. This is consistent with the evidence of material culture exchange between Sardinians and mainland Mediterranean groups^15^, although genome-wide analyses find no significant evidence of Steppe ancestry so the quantitative demographic impact was minimal. *qpAdm* modeling of the ancestry of the *Sardinia_Nuragic_BA10365* outlier with respect to sources potentially more closely related in space and time does infer some ancestry in this individual from an eastern source (either carrying Steppe ancestry or Iranian-related ancestry) that we do not detect by modeling with sources more distant in space and time, consistent with the hypothesis of eastern influence (**Supplementary Table 12**).

We detect definitive evidence of Iranian-related ancestry in an Iron Age Sardinian I10366 (391-209 calBCE) with an estimate of 11.9 ± 3.7.% *Iran_Ganj_Dareh_Neolithic* related ancestry, while rejecting the model with only *Anatolian_Neolithic* and *WHG* at p=0.0066 (**Supplementary Table 9**). The only model that we can fit for this individual using a pair of populations that are closer in time is as a mixture of *Iberia_Chalcolithic* (11.9 ± 3.2%) and *Mycenaean* (88.1 ± 3.2%) (p=0.067). This model fits even when including Nuragic Sardinians in the outgroups of the *qpAdm* analysis, which is consistent with the jhypothesis that this individual had little if any ancestry from earlier Sardinians.

In the *Sardinian_LateAntiquity* group (the earliest dating to 256-403 calCE), we detect even higher proportions of *Iran_Ganj_Dareh_Neolithic*-related ancestry: an estimated 29.6 ± 4.6.% (p=0.000001 for rejection of the alternative model that attempts to model its eastern ancestry as entirely Yamnaya-related, **Supplementary Table 9**). One possibility is the Iranian-related ancestry began to be introduced in the Phoenician period, a scenario that is not only consistent with the historical evidence and our finding of this ancestry type in the Iron Age Sardinian, but is also supported by previously published mitochondrial DNA which has documented haplotypes in ancient Phoenician colonies in modern Sardinians^51^. In modeling using source populations that are temporally more plausible, this individual is consisten with being a clade with both *Myceanean* (p=0.241) or *Ibiza_Phoenician* (p=0.145); importantly, both these models works with Nuragic Bronze Age Sardinians included in the outgroups, and so *Sardinian_LateAntiquity* is consistent with having negligible ancestry from earlier Bronze Age groups to the limits of our resolution (**Supplementary Materials**). We also model the outlier *Sardinia_LateAntiquity12221* as having 33.3 ± 5.5% Yamnaya-related while confidently rejecting models with no Steppe ancestry (all p≤0.001) (**Supplementary Table 9**), providing the earliest clear evidence of Steppe ancestry in Sardinia. However, we do not have sufficient resolution given the limited data from this single sample to determine the geographic source of the Steppe ancestry (**Supplementary Table 13**).

In a dataset of 27 modern Sardinians for whom we have genotyping data at about 600,000 SNPs^45^, we obtain a fit for a model of 61.4 ± 1.6% *Anatolia_Neolithic,* 9.5 ± 1.0% *WHG,* 19.1 ± 1.9% *Iran_Ganj_Dareh_Neolithic* and 10.0 ± 1.6% *Yamnaya_Samara* related ancestry and definitively reject models without all four ancestries (all models p<10^-6^ in **Supplementary Table 9**). We replicate the finding of *Iran_Ganj_Dareh_Neolithic*-related ancestry (and not just Steppe ancestry) in a subset of four of the modern Sardinian individuals with whole genome shotgun sequencing data (**Supplementary Table 9**). Even the four-way model is not comprehensive for modern Sardinians, however, as when we add Late Neolithic North Africans from Morocco to the outgroup set^52^, we reject the four-way mixture model (p<10-^12^) (adding the Neolithic Moroccans to the outgroup set does not cause model rejection for any of the ancient samples in our dataset, showing that it may reflect events taking place after the times our individuals lived; **Supplementary Table 9**). Modeling modern Sardinians with this fifth sources produces a fit with an estimate of 16.1 ± 8.4% *Morocco_LN*-related ancestry (p=0.235). Our signal of North African-related mixture in Sardinians may reflect the same process that introduced sub-Saharan African ancestry into Sardinians^53–55^ which was argued in ^56^ to reflect North African-related admixture with an average date of ∼630 CE.

An important question is how much ancestry modern Sardinians have inherited from people related to those of the Nuragic Bronze Age. We could parsimoniously model our modern Sardinian sample as a 2-way mixture of 13.6 ± 3.4% *Sardinia_Nuragic_BA* and 86.4 ± 3.4% *Sardinia_LateAntiquity12221*. It is striking that most of the ancestry in modern Sardinians is inferred in this analysis to come from a *Sardinia_LateAntiquity12221*-related group, which can itself be modeled as closely related to Mycenaeans or Phoenicians with no evidence of specific shared ancestry with Bronze Age Sardinians. The group of modern Sardinians we are modeling has often been interpreted as an isolated lineage that derives from early Sardinian farmers with little subsequent immigration into the islands. Our finding that a large fraction of this group’s ancestry is consistent with deriving from a group that was present in Sardinia in Late Antiquity and that had no evidence of a contribution from earlier Sardinian groups is therefore surprising (although we caution that this inference is tentative as the amount of data we have for *Sardinia_LateAntiquity12221* is limited; **Online Table 1**). Modern Sardinian populations are geographically highly substructured for example among different valleys and coastal and inland sites.^55^ Analyses of more geographically diverse modern and ancient Sardinians will provide additional insight into the population turnovers.

### Formal Modeling of the Neolithic to Bronze Age transition in Sicily

In the Middle Neolithic, Sicilians harbored ancestry typical of early European farmers, well modeled as a mixture of *Anatolia_Neolithic* and *WHG* (**Fig. 2, Fig. 4, Supplementary Table 9**).

Steppe ancestry arrived in Sicily by the Early Bronze Age. While a previously reported Bell Beaker culture-associated individual from Sicily had no evidence of Steppe ancestry^4^, a result we confirm by more than tripling the number of sequences for this individual who previously had marginal quality data, we find evidence of Steppe ancestry in the Early Bronze Age by ∼2200 BCE. In distal *qpAdm*, the outlier *Sicily_EBA11443* is parsimoniously modeled as harboring 40.2 ± 3.5% Steppe ancestry, and the outlier *Sicily_EBA8561* is parsimoniously modeled as harboring 23.3 ± 3.5% Steppe ancestry (**Fig. 4a, Supplementary Table 9**). The main *Sicily_EBA* cluster also can only be fit with Steppe ancestry albeit at a lower proportion of 9.1 ± 2.3%, and models without Steppe ancestry can be rejected (p=0.001) (**Supplementary Table 9**). The presence of Steppe ancestry in Early Bronze Age Sicily is also evident in Y chromosome analysis, which reveals that 4 of the 5 Early Bronze Age males had Steppe-associated Y-haplogroup R1b1a1a2a1a2. (**Online Table 1**). Two of these were Y-haplogroup R1b1a1a2a1a2a1 (Z195) which today is largely restricted to Iberia and has been hypothesized to have originated there 2500-2000 BCE^57^. This evidence of west-to-east gene flow from Iberia is also suggested by *qpAdm* modeling where the only parsimonious proximate source for the Steppe ancestry we found in the main *Sicily_EBA* cluster is Iberians (**Supplementary Table 14**).

We detect Iranian-related ancestry in Sicily by the Middle Bronze Age 1800-1500 BCE, consistent with the directional shift of these individuals toward Mycenaeans in PCA (**Fig. 2b**). Specifically, two of the Middle Bronze Age individuals can only be fit with models that in addition to *Anatolia_Neolithic* and *WHG*, include *Iran_Ganj_Dareh_Neolithic*. The most parsimonious model for *Sicily_MBA3125* has 18.0 ± 3.6% Iranian-related ancestry (p=0.032 for rejecting the alternative model of Steppe rather than Iranian-related ancestry), and the most parsimonious model for *Sicily_MBA4109* has 14.9 ± 3.9% Iranian-related ancestry (p=0.037 for rejecting the alternative model) (**Fig. 4a, Supplementary Table 9**). This inference is also supported by *qpAdm* using sources closer in geography and time that always identify a parsimonious model with *Minoan_Lassithi* as a source for these two individuals (**Supplementary Table 15**). We also found evidence of Iranian-related ancestry in Sicily in an individual of the Early Bronze Age cluster, I11442, who could only be fit in a 3-way model with Iranian-related ancestry (19.3 ± 3.8% ancestry of this type, p=0.391; the 3-way model involving Steppe ancestry fails to a fit (p=0.010)) (**Supplementary Table 10**). However, this finding should be viewed with caution as *qpWave* clustered this individual with four other Sicilian Early Bronze Age individuals, so this finding could be an artifact of performing tests on our data beyond what is justified by our groupings. The modern southern Italian Caucasus-related signal identified in ^58^ is plausibly related to the same Iranian-related spread of ancestry into Sicily that we observe in the Middle Bronze Age (and possibly the Early Bronze Age).

For the Late Bronze Age group of individuals, *qpAdm* documented Steppe-related ancestry, modeling this group as 80.2 ± 1.8% *Anatolia_Neolithic*, 5.3 ± 1.6% *WHG*, and 14.5 ± 2.2% *Yamnaya_Samara* (**Fig. 4b, Supplementary Table 9**). Our modeling using sources more closely related in space and time also supports *Sicily_LBA* having Minoan-related ancestry or being derived from local preceding populations or individuals with ancestries similar to those of *Sicily_EBA3123* (p=0.527), *Sicily_MBA3124* (p=0.352), and *Sicily_MBA3125* (p=0.095) (**Supplementary Table 15**).

Finally, when we model modern Sicilians, we find that they require not only Steppe and Iranian-related ancestries but also North African ancestry, confirming the ample historical and archaeological evidence of major cultural impacts on the island from North Africa after the Bronze Age (**Supplementary Materials**).

## Discussion

The islands of the western Mediterranean have been among the most poorly studied regions of Europe from the perspective of genome-wide ancient DNA. Here we increase by about 17-fold the number of individuals with data from the Neolithic onward in these islands to document the arrival of both Steppe and Iranian-related ancestry.

In the Balearic islands, we show that Steppe ancestry arrived almost simultaneously with the first permanent human occupation of the islands in the Early Bronze Age, while the North African ancestry that arrived at least by the time of the Phoenicians^50^ still is present today. In Sicily, Steppe ancestry arrived by ∼2200 BCE, and likely came at least in part from the west as it was associated with the Iberian-specific Y haplogroup R1b1a1a2a1a2a1 (Z195),^57^ thus documenting how Iberia was not just a destination of east-to-west human movement in Europe, but also an important source for west-to-east Steppe ancestry reflux^59^. In Sardinia, we find no convincing evidence of Steppe ancestry in the Bronze Age, but we detect it by ∼200-700 CE.

We find no evidence of Iranian-related ancestry in the Balearic Islands individuals until the Phoenician period, around the same time as we detect it in Sardinia. In Sicily, Iranian-related ancestry was present during the Middle Bronze Age, showing that this ancestry which was widespread in the Aegean around this time (in association with the Minoan and Mycenaean cultures), also reached further west. Based on our analysis of modern individuals, it is possible that this ancestry first spread west in substantial amounts during the Late Helladic period of the Mycenaean expansion when strong cultural interactions between Sicily and the Aegean are documented^18,60–62^. However, if our signal of such ancestry in an Early Bronze Age Sicilian individual is correct then some of this spread began even earlier.

Our co-analysis of modern and ancient Sardinians questions the commonly held view that Sardinians are well described as an isolated remnant of Europe’s first farmers^63^. While Nuragic Bronze Age Sardinians are indeed well-modeled as having a typical early European farmer ancestry profile, modern Sardinians harbor substantial fractions of ancestry from several groups that arrived in Europe after the Neolithic, and we model modern Sardinians as harboring 10.0 ± 1.6% Steppe ancestry and an even larger 19.1 ± 1.9% Iranian-related ancestry. Both ancestry types are definitively required to model modern Sardinians, and we show that modern Sardinians have been substantially impacted by movement of ancestry from North Africa in the last two millennia. Thus, rather than being an island sheltered from admixture and migration since the Neolithic, Sardinia, like almost all other regions in Europe has, been a site for major movement and mixtures of people.

## Materials and Methods

### Laboratory work details

We ground skeletal samples to powder in dedicated ancient DNA facilities at the University College Dublin in Ireland, at the University of Florence in Italy, at the University of Palermo in Italy, and at Harvard Medical School in Boston USA (**Online Table 1**)^22,64,65^. We treated all DNA extracts with Uracil-DNA Glycosylase (UDG) to remove characteristic ancient DNA damage to cleave the molecules at 5’ Uracils, thus reducing the rate of damage-induced errors^23^. For two of the samples, we performed DNA extraction^19,20^ and double-indexed library preparation in Florence^23^. For all other samples, we performed DNA extraction at Harvard Medical School, sometimes using silica coated magnetic beads to support robotic cleanups (instead of silica column cleanups that were used for manual DNA extraction)^19,21^. We converted these DNA extracts to individually barcoded libraries, in some cases assisted by a robotic liquid handler^23^ (see **Online Table 1** for details). We initially screened libraries by enriching the libraries for the human mitochondrial genome^66^ and about 3000 nuclear SNPs using synthesized baits (CustomArray Inc.), and sequencing on an Illumina NextSeq500 instrument, using different index pairs to distinguish between them. We merged read pairs that overlapped by at least 15 base pairs allowing up to one mismatch (and representing each overlapping base by the higher quality base), and computationally trimmed adapters and barcodes. We mapped the merged sequences to the reconstructed human mitochondrial DNA consensus sequence^67^ using bwa (v.0.6.1)^68^, and removed duplicate sequences that had the same orientation, same start and stop positions, and the same barcodes. We assessed the data for authenticity by computing the damage rate at the terminal cytosines (which we required to be at least 3% for at least one library for each individual following published recommendations for libraries of this type^23^), and by estimating the rate of mismatches to the consensus mitochondrial sequence using contamMix^24^ We next enriched the samples with promising quality for 1233013 SNPs (‘1240K SNP capture’)^2,25^, and sequenced and processed them as for the mitochondrial DNA with the difference that we mapped to the human reference genome *hg19*. We assessed authenticity as for the mitochondrial DNA data, while also estimating contamination based on the ratio of Y to X chromosome sequences (filtering out individuals that had a ratio unexpected for a male or a female) as well as the rate of heterozygosity at X-chromosome positions (only valid as an estimate of contamination in males who should have no X chromosome variation^69^. For some libraries we co-enriched samples for the mitochondrial genome together with the 1240k targets (“1240k+” enrichment).

### Radiocarbon dating and quality assurance

We performed 25 accelerator mass spectrometry (AMS) radiocarbon dates (14C) on samples from 24 skeletons at the Pennsylvania State University (PSU) Radiocarbon Laboratory, as well as an additional 4 direct dates on an additional 3 samples. Here we give a detailed description of the samples processing at PSU, as it is the source of most of our dates (for the other samples, we refer readers to the published protocols). As precaution at PSU, we removed possible contaminants (convervants/adhesives) by sonicating all bone samples in successive washes of ACS grade methanol, acetone, and dichloromethane for 30 minutes each at room temperature, followed by three washes in Nanopure water to rinse. We extracted bone collagen and purified using a modified Longin method with ultrafiltration (>30kDa gelatin^70^). If collagen yields were low and amino acids poorly preserved we used a modified XAD process (XAD Amino Acids^71^). For quality assurance, we measured carbon and nitrogen concentrations and C/N ratios of all extracted and purified collagen/amino acid samples with a Costech elemental analyzer (ECS 4010). We evaluated sample quality by % crude gelatin yield, %C, %N and C/N ratios before AMS 14C dating. C/N ratios for all directly radiocarbon samples fell between 2.9 and 3.6, indicating excellent preservation^72^. We combusted collagen/amino acid samples (∼2.1 mg) for 3 h at 900°C in vacuum-sealed quartz tubes with CuO and Ag wires. Sample CO2 was reduced to graphite at 550°C using H2 and a Fe catalyst, and drew off reaction water with Mg(ClO4)2^73^. We pressed graphite samples into targets in Al boats and loaded them onto a target wheel with OX-1 (oxalic acid) standards, known-age bone secondaries, and a 14C-free Pleistocene whale blank. We made all 14C measurements on a modified National Electronics Corporation compact spectrometer with a 0.5 MV accelerator (NEC 1.5SDH-1). We corrected the 14C ages for mass-dependent fractionation with measured δ13C values^74^ and compared with samples of Pleistocene whale bone (backgrounds, 48,000 14C BP), late Holocene bison bone (∼1,850 14C BP), late 1800s CE cow bone, and OX-2 oxalic acid standards. We calibrated 14C ages with OxCal version 4.3^75^ and the IntCal13 northern hemisphere curve^76^. The stable carbon and nitrogen isotope measurements we obtained do not indicate a large marine dietary component in these individuals despite their coming from island populations and hence we did not perform a correction of the dates for marine reservoir effect.

### Uniparental haplogroup determination

We determined mitochondrial haplogroups using HaploGrep^77^ and phylotree^78^ (build 17) on the data from the mitochondrial enrichment experiment^79^. We restricted sequences and base qualities to values of ≥30, and built a consensus sequence with *samtools* and *bcftools*^*80*^, using a majority rule and minimum coverage of 1, trimming 2 basepairs from the end of each sequence. We further restricted the data for each sample to the damaged reads as determined by *pmdtools* (using a minimum *pmdscore* of 3) and repeated the calling. In almost every case where there was sufficient post-damage restricted coverage to give a confident haplogroup call, the calls matched the non-restricted read sample. We restricted sequences for Y-chromosome haplogroup assessment to qualities ≥30, and identified the most derived mutations using the nomenclature of the International Society of Genetic Genealogy (http://www.isogg.org) version 11.110.

### Dataset assembly

We assembled a base dataset and then subsetted for each analysis. This complete dataset included 3310 individuals, of which 2191 were modern^27,39,45–47^ and 1119 were ancient individuals from previous publications ^1,2,4,6,26–44,52^, which we combined with the newly reported 49 samples (**Online Table 3**). We performed all subsequent analysis on autosomal data.

### Principal component analysis

We used a subset of 736 modern and 1123 ancient West Eurasians for principal component analysis (PCA) using *smartpca* from the EIGENSOFT package^81^. We modified the standard parameter file with the options shrinkmode: YES, and lsqproject: YES to project all ancient individuals onto the eigenvectors computed from modern vectors. We used a dataset containing only transversions to assess the robustness of our qualitative inferences to bias due to ancient DNA damage-induced errors (**Supplementary Fig. 1**).

### Population structure analysis

We ran ADMIXTURE^48^ after pruning to remove one SNP each in pairs of SNPs in linkage disequilibrium, using PLINK1.9^82^ and the option --indep-pairwise 200 25 0.4, leaving 321518 SNPs. We ran ADMIXTURE from K=5 to K=15, with 5 random-seeded replicates for each value of K. We used cross validation by adding the option --cv to find the runs with the lowest errors. For each value of K, we kept the replicate with lowest error. We present results for K=10, as we empirically found that this is the value of K with lowest cross-validation error that also showed clear distinctions between ancient Western, Eastern, and Caucasus Hunter-Gatherer backgrounds, while having a maximized Early Neolithic Anatolian component. We also performed ADMIXTURE restricting to transversion SNPs and obtained qualitatively similar results suggesting that ancient DNA damage is unlikely to be strongly biasing our findings (**Supplementary Fig. 1**).

### *f*_4_-statistics

We used ADMIXTOOLS^45^ to compute *f*_*4*_-statistics (*qpDstat*). We used *Mbuti.DG* as our outgroup, and computed statistics of the form *f*_*4*_(*Mbuti.DG*, X; Y, Z), where X is our test population/individual and Y/Z are pairs to test against. We used the options f4mode: YES and printsd: YES. We used *f*_*4*_-statistics to assess overall population affinities and changes in ancestry through time either by direct comparison of the test populations with the desired pairs or by using symmetry tests, where the populations Y and Z are the populations being tested for consistent with descent from a common ancestral population.

### *qpWave*/*qpAdm*

We used *qpWave*/*qpAdm* from ADMIXTOOLS^45^ to estimate admixture coefficients and to model our individuals/populations as result of groups related to different proxies for the true source population. We used a base outgroup set including the following individuals/populations: *Mbuti.DG, Ust_Ishim, CHG, EHG, ElMiron, Vestonice16, MA1, Israel_Natufian, Jordan_PPNB*. Extra populations were included in each test to improve accuracy when using populations with similar ancestries (see **Supplementary Materials** for a detailed description). When analyzing the results we present the most parsimonious model with the highest probability. We used the option allsnps: YES.

## Data Availability

All raw data are available at the European Nucleotide Archive and the National Center for Biotechnology Information under the accession number [to be included upon paper acceptance] and at https://reich.hms.harvard.edu/datasets.

## Acknowledgements

This manuscript is dedicated to the memory of Sebastiano Tusa of the Soprintendenza del Mare in Palermo, who would have been an author of this study had he not tragically died in the crash of Ethiopia Airlines flight 302 on March 10. We thank Zhao Zhang for database support. We thank the Soprintendenza BBCCAA Palermo and Rosario Schicchi (director of Museum of Castelbuono) for facilitating access to important skeletal materials. D.F. was supported by an Irish Research Council grant GOIPG/2013/36. Radiocarbon work was supported in part by the NSF Archaeometry program BCS-1460369 to D.J.K. and B.J.C. C.L.-F. was supported by Obra Social La Caixa and by FEDER-MINECO (BFU2015-64699-P). D.C. was supported by the grant 20177PJ9XF MIUR PRIN 2017. D.Re. is an Investigator of the Howard Hughes Medical Institute and his ancient DNA laboratory work was supported by National Science Foundation HOMINID grant BCS-1032255, by National Institutes of Health grant GM100233, by an Allen Discovery Center grant, and by grant 61220 from the John Templeton Foundation.

## Author Contributions

D.M.F., D.Re., and R.P. conceived the study. D.M.F., E.C., C.C., G.C., M.C., V.F., M.Lo., E.M., Ma.M., R.M.M., D.Ra., M.R.P., V.S., P.S., L.T, M.T-N., C.L-F, L.S., D.C., R.P. excavated, assembled and/or studied the osteological material. D.M.F., O.C., N.R., N.B., M.F., B.G., M.La., Me.M., A.Mo., M.N., J.O., K.A.S., K.S., and S.V. performed laboratory work, while N.R., D.C., and R.P. supervised this work. J.C. provided computing resources. B.J.C. performed radiocarbon analysis and D.J.K. supervised this work. D.M.F., I.O., R.B., S.M., and M.Ma. performed bioinformatic and population genetic analysis with input from A.Mi., I.L., N.P., and D.R.

