## Supplemental_Materials for "The Arrival of Steppe and Iranian Related Ancestry in the Islands of the Western Mediterranean"

### SUPPLEMENTARY FIGURES

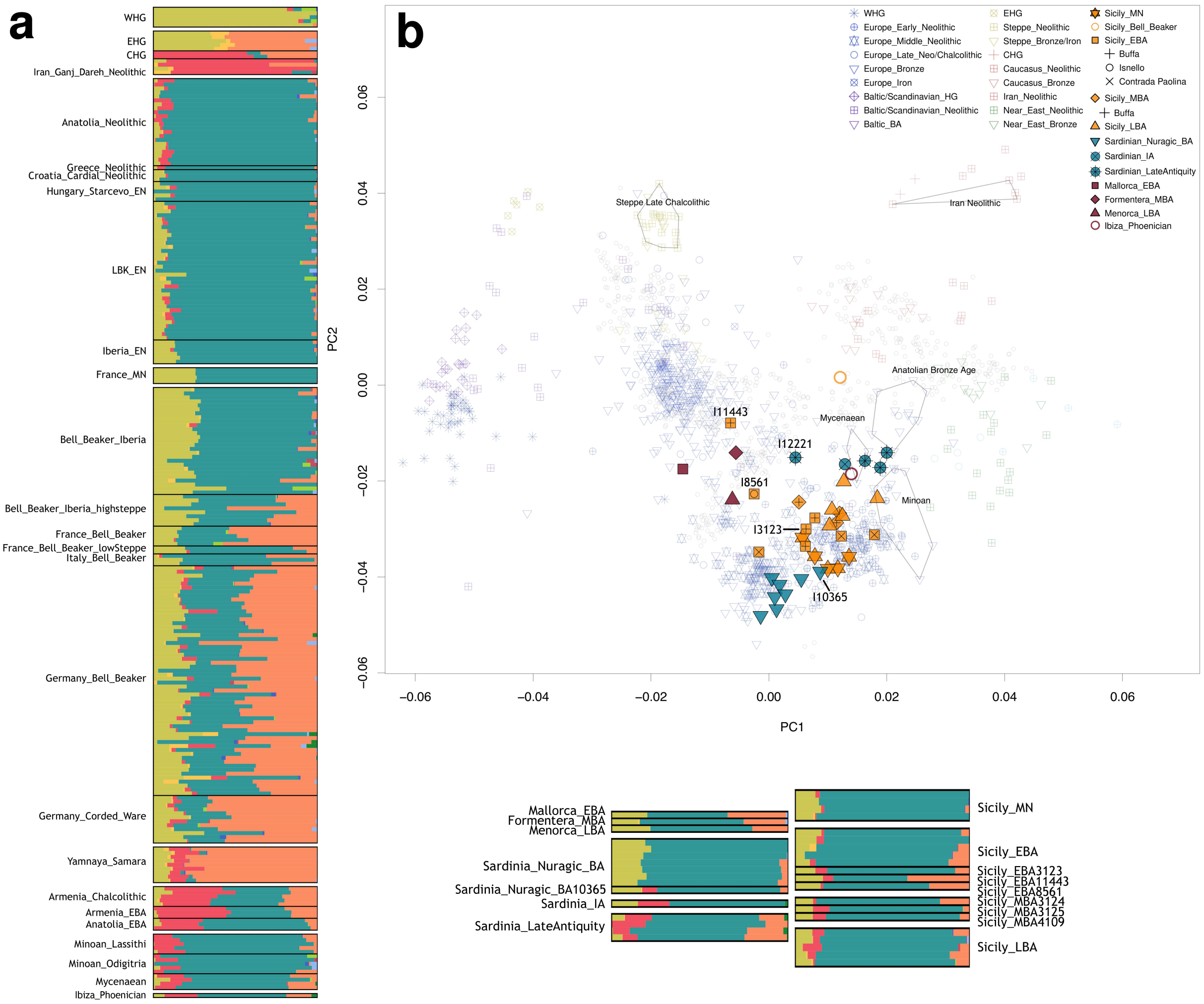


**Supplementary Fig. 1: Qualitative relationships between ancient Sardinians, Sicilians and Balearic islanders and other ancient and present-day populations using a dataset restricted to only transversion SNPs according to a) unsupervised cluster-based ADMIXTURE analysis with K=10; and b) PCA with previously published ancient individuals (non-filled symbols), projected onto variation from present-day populations (gray squares).**

**SUPPLEMENTARY TABLES**

**(as Excel file) Online Table 1.** Sample information, sequencing, and contamination results for the new individuals from this study.

**(as Excel file) Online Table 2.** Information about literature samples used in this study.

**(as Excel file) Online Table 3.** New direct radiocarbon dates presented in this study.

#### SUPPLEMENTARY NOTES

##### *1 - Archaeological descriptions and contexts of newly reported samples*

*Cova des Moro, Mallorca, Spain ( ⊕ 39.505, 3.302)*

Cova des Moro is a natural cave in East Mallorca (municipality of Manacor) with a paleontological and archaeological deposit. It is found in a Miocene coastal cliff and formed by a large hall (about 60 by 30 m) divided into different spaces by stalagmite formations[1]. Archaeological excavations took place between 1995 and 2002[2,3], identifying three main periods of occupation. The earliest was dated to the second half of the 3rd millennium BCE. The material evidence (pottery and animal bones) indicates that in this period it was used as a temporary shelter. The second period, dated to the Late Bronze Age (mid and late 2nd millennium BCE), has been interpreted as a ritual use of the cave, during which a cyclopean corridor at the entrance of the cave was probably built. Lastly, the cave was used as refuge by Muslims during the Catalan conquest of Mallorca (1229-1232 CE).

Far from the entrance, several human bones were found in a shallow sinkhole in the central part of the cave. Radiocarbon dates were determined on two human samples, UtC-7878: uncal3840±60 uncalBP[4], and KIA-30020: uncal3900±30 BP[5], relating these human bones to the early occupation period of the cave. The following sample from this site was used (same format for sample description used for all sites below):

**CDMDR (*Mallorca_EBA*, I4329).** Various skull and mandible fragments that probably belonged to the same individual. C14 dated to 2395-2316 calBCE (3900±30 BP, KIA-30020)

*Cova 127, Formentera, Spain ( ⊕ 38.670, 1.584)*

Cova 127 is a cave located om the cliffs at La Mola (Formentera) and was excavated from 2014 to 2016. The cave entrance connects to a gallery that leads to a main hall. A burial with two skeletons was found in the junction between both areas, consisting of fragmented remains from two individuals[6]: one middle-aged adult around 35 years old at death (used in this study) and one juvenile of 10-15 years old. Some bone buttons were recovered (one with a V-shaped perforation) and ceramic fragments were uncovered along with the human remains. The ascription to the Bronze Age period is supported not only by the direct radiocarbon dating of the human bones, but also by cultural similarities with the early occupation layers of the nearby Cap de Barbaria II settlement that is currently being excavated[6].

**Cova127, 2015, Ni 1077; No. Estracio 5209 (*Formentera_MBA*, I4420)**. Petrous bone, from an assemblage of fragmented human bones (n=159) and isolated teeth from two individuals from Cova 127. We newly report two C14 dates for I4420 - a skull piece dated to 1879-1691 calBCE

(3454±26 BP, D-AMS018425), and a maxillar piece dated to 1881-1701 calBCE (3473±20 BP, MAMS-22647). We use the union of these date ranges, 1881-1961 calBCE for I4420, to represent I4420.

**Cova127, 2015, Ni 1083, No. Estacio 5202(CR), UE 1 (*Formentera_MBA*, I8573)**. Tooth, from an assemblage of fragmented human bones (n=159) and isolated teeth from two individuals from Cova 127. For this individual, we generated a date based an associated (probably same individual) bone from the site with a date of 1883-1695 calBCE (3469±28 BP, D-AMS-018426).

*Naveta des Tudons, Menorca, Spain (⊕ 40.003, 3.892)*

This funerary Cyclopean building in West Menorca (municipality of Ciutadella) has a maximum length of 13.6 m, maximum width of 6.4 m, and height of 4.5 m. During the excavations 1959-60 while it was being restored, human remains were recovered from a collective burial. Four radiocarbon dates were determined on the human samples - IRPA-1178: uncal2740±40 BP, IRPA-1181: uncal2780±35 BP, IRPA-1184: uncal2690±35 BP, uncalIRPA-1179: uncal2820±40 BP[7]. These dates all fall in between ca. 1100-800 calBCE, and cluster in the first half of the 9th century calBCE. This chronology, as well as the materials recovered, place this individual into a Talaiotic context. "Naveta" is a Catalan word which means small boat, and is locally used to name these buildings because of their similarity with inverted boats. The burial naveta, as a type of funerary monument, is considered to be a local development (only documented in Menorca) derived from the dolmens of the Early Bronze Age[8,9]. The elongated and stylized construction of Naveta des Tudons probably represents the final evolution of this type of burial structure.

**NT14 (*Menorca_LBA*, I3315)**. Recovered from an assemblage of at least 100 individuals in a collective inhumation[10]. C14 dated to 904-817 calBCE (2715±20 BP, PSUAMS-3717).

*Stretto Partanna, Sicily, Italy (⊕ 37.724, 12.916)*

The excavation at the ditch-trench of Stretto Partanna (Trapani) took place in 1989. The archaeological materials consist predominantly of trichromatic ceramics, and radiocarbon dates attributed the occupation period to the Middle Neolithic (5800-4900 BCE)[11]. The ditch-trench, over 13 meters deep, was characterized by the presence of artifacts along with skeletal remains of domesticated fauna. The skeletal human material collected consisted of at least 7 secondary burials that were not anatomically connected [11]. The data in this study came from 4 petrous bones:

**FS2 (*Sicily_MN*, I4062).** C14 dated to 4946-4787 calBCE (5980±30 BP, PSUAMS-1950).

**FS3 (*Sicily_MN*, I4063).** C14 dated to 4963-4795 calBCE (5995±30 BP, PSUAMS-2263).

**FS4 (*Sicily_MN*, I4064).** C14 dated to 4837-4713 calBCE (5900±30 BP, PSUAMS-2266).

**FS5 (*Sicily_MN*, I4065).** C14 dated to 4975-4781 calBCE (5980±35 BP, PSUAMS-1951).

<u>*Buffa Cave II, Sicily, Italy (⊕ 37.908, 13.478)*</u>

The human skeletal remains from the cave of Buffa II (near Villafrati, a comune in the Province of Palermo, located about 25 km southeast of Palermo), were recovered on behalf of Ferdinand Freiherr von Andrian-Werburg (1835-1913) in the winter of 1876/77. Andrian-Werburg entrusted Domenico Heina, servant in the Geologic Cabinet of the University of Palermo and collector of natural produce, to carry out the excavations. Besides a mix of different archaeological objects and animal remains, he recovered human cranial and postcranial remains of a minimum of 21 individuals. Andrian-Werburg published the results in 1878 in the article "Prähistorische Studien aus Sicilien" in a Supplement booklet of volume X of the Berliner Zeitschrift für Ethnologie. His descriptions are divided into two parts: the first chapter deals with Palaeolithic findings, and the second with Neolithic/Chalcolithic[12].

More recent work on Buffa Cave II identified an occupation extending into the Early Bronze Age, as evidenced by the finding of pottery belonging to the Bell Beaker and Capo Graziano cultures[13]. Von Andrian-Werburg's study analyzes the human remains initially attributed to the Neolithic/Chalcolithic. The cranial and postcranial remains were investigated by Emil Zuckerkandl, prosector at the Institute of Anatomy, and then handed over as a "personal present" to the newly founded Anthropological Ethnographical Department at the Natural History Museum Vienna. The cranial and postcranial elements were inventoried 13th December 1890 by C. Heinzel und J. Szombathy with the numbers 3029 till 3062 (a total of 179 bone pieces). The remains belong most likely to the Neolithic ("Neolithische Schichte"), although Andrian-Werburg recognised cultural differences with phases that he could not clearly separate[12]. Direct radiocarbon analysis of three of the individuals identified the remains as belonging to the Early and Middle Bronze Age, closer to what is described in [13].

The following details are extracted from the handwritten notes (Inventory register no. 3, pages 00027-00029) and Zuckerkandl[14], who concentrated in his comprehensive study on the anatomical and morphological features of the specimens from Buffa cave near Villafrati.

**BU32, grave 3032 (*Sicily_EBA11443*, I11443).** Petrous bone. C14 dated to 2279-2102 calBCE (4090±60 BP, OxA-32773).

**BU30, grave 3030, cranium II (*Sicily_EBA*, I3122).** Skull, with portions of the cranial base missing, slightly asymmetric, with worn teeth and some cavities. The skull sutures are open and the age at death is estimated at ca. 25-30 years. Morphologically the individual is inferred to be a male which is contradicted by the genetic sex determination of female. C14 dated to 2266-2032 calBCE (3730±30 BP, PSUAMS-1985).

**BU36C, grave 3036 (*Sicily_EBA*, I11442).** One of four temporal bones preserved from grave 3036 (2 from the left and 2 from the right side). C14 dated to 2281-2047 calBCE based on a union of two radiocarbon dates [2273-2047 calBCE (3750±20 BP, PSUAMS-4547), 2281-2062 calBCE (3765±20 BP, PSUAMS-4620)].

**BU31, grave 3031, cranium III (*Sicily_EBA3123*, I3123).** Skull, with the left portion of the frontal bone missing, extremely brachycephalic ("planoccipital"). The skull sutures are open. Morphologically the individual is inferred to be a male. C14 dated to 2287-2044 calBCE (3760±30 BP, PSUAMS-3892).

**BU36, grave 3036 (*Sicily_MBA*, I3124).** One of four temporal bones preserved from grave 3036 (2 from the left and 2 from the right side). C14 dated to 1948-1777 calBCE (3545±20 BP, PSUAMS-4317).

**BU36A, grave 3036 (*Sicily_MBA*, I3125).** One of four temporal bones preserved from grave 3036 (2 from the left and 2 from the right side). C14 dated to 1612-1506 calBCE (3275±20 BP, PSUAMS-4318).

**BU36B, grave 3036 (*Sicily_MBA*, I4109).** One of four temporal bones preserved from grave 3036 (2 from the left and 2 from the right side). C14 dated to 1631-1509 calBCE (3300±25 BP, PSUAMS-1949).

*Marcita, Sicily, Italy ( ⊕ 37.680, 12.789)*

The Marcita necropolis (Castelvetrano - Trapani) was dated between the Early and the Middle Bronze Age. The excavations took place in 1984, when three rock-cut tombs were revealed: A, B and C. Tombs A and B included archaeological finds attributed to the Bell Beaker culture, while Tomb C had human remains but almost no archaeological assemblage (only an ivory comb)[15]. It was composed of about 70 individuals in secondary deposition and not in anatomical connection. The individuals seem to have been placed randomly and all together, perhaps related to a violent event[15].

**MA4 (*Sicily_LBA*, S10373.E1.L2).** Dated to C14 dated to 1450-1150 BCE based on being the son of I3878 at 1379-1196 calBCE (3015±20 BP, PSUAMS-3995)].

**MA8 (*Sicily_LBA*, I3878_new).** C14 dated to 1379-1196 calBCE (3015±20 BP, PSUAMS-3995).

**MA1 (*Sicily_LBA*, S10371.E1.L2).** Context dated to 1400-900 BCE based on C14 dates from other individuals at the same site.

**MA2 (*Sicily_LBA*, S10372.E1.L2).** Context dated to 1400-900 BCE based on C14 dates from other individuals at the same site.

**MA3 (*Sicily_LBA*, I3876).** C14 dated to 1071-902 calBCE (PSUAMS-1986).

*Alghero, Sardinia, Italy ( ⊕ 40.55, 8.31)*

Lu Maccioni (Alghero, Province of Sassari) is a natural cave located in Northern Sardinia, precisely in the historical-geographical region of Nurra, at sea level (Province of Sassari). Pieces of dark ceramic and a cylinder in white marble have been found in the cave. The 19 skulls and the 57 long bones recovered allowed the estimation of a NMI of 40 individuals. Teeth showed presence of caries and traces of trauma have been observed in the skulls.

Relative dating and radiocarbon analysis traced back the site to final Neolithic. The site originates during the Ozieri culture (late Neolithic), but it was reused during the Nuragic III (Bronze Age) and the Phoenician period (Iron Age).

The osteological collection of Lu Maccioni is housed in the Sardinian Museum of Anthropology and Ethnography of the University of Cagliari[16].

**Lu Maccioni_Ind.A (*Sardinia_Nuragic_BA*, I10364).** *Petrous bone,* context dated to 1150-800 BCE based on a C14 date of 1126-825 calBCE (2810±60 BP, Beta-82329) from the same layer.

**Sardinia, Lu Maccioni_31 Neo-Iron_23 (*Sardinia_Nuragic_BA*, I3642).** Tooth, C14 dated to 1117-976 calBCE (2870±20 BP, PSUAMS-2387).

*Perdasdefogu, Sardinia, Italy ( ⊕ 39.683, 9.433)*

Perdasdefogu cave is a natural cave located in the historical-geographical region of Quirra, central eastern Sardinia, at 535 m above sea level. It consists of a single corridor, about 605 m long, 90 cm wide, and with a height varying from 0.70-2 meters.

The excavations took place in 1963. The cave had been damaged by human action. No funerary utensils, ornaments, or other archaeological materials were found. Due to the collective burial ritual, no articulated skeletons were found, but several human bones, including 36 skulls and 58 long bones, were collected and attribute to a minimum number of 50 individuals.

The ascription to the final Bronze Age period is supported by radiocarbon dating.

The osteological collection of Perdasdefogu cave is housed in the Sardinian Museum of Anthropology and Ethnography of the University of Cagliari[16].

**Perda-1 (*Sardinia_Nuragic_BA*, S10552.E1.L1).** Petrous bone, C14 dated to 1384-1213 calBCE (3025±20 BP, PSUAMS-4875).

**Perda-2 (*Sardinia_Nuragic_BA*, S10553.E1.L1).** Petrous bone, C14 dated to 1226-1056 calBCE (2950±20 BP, PSUAMS-4876).

**Perda-3 (*Sardinia_Nuragic_BA*, S10554.E1.L1).** Petrous bone, C14 dated to 1260-1112 calBCE (2960±20 BP, PSUAMS-4877).

**PERDASDEFOGU 15R (*Sardinia_Nuragic_BA*, I3741).** Tooth, C14 dated to 1219-1049 calBCE (2935±25 BP, PSUAMS-2118).

*Seulo, Sardinia, Italy ( ⊕ 39.867, 9.233)*

Stampu Erdi (Seulo) is a natural cave located in central southern Sardinia, in the historical-geographical region of Barbagia of Seulo, at 740 m above sea level. Archaeological excavation took place in 1956. Besides human cranial and postcranial remains, pottery and objects in obsidian have been found. The site was dated back at Bronze age (Nuragic II period) through both by relative and by radiocarbon analysis.

The osteological collection of Stampu Erdi is housed in the Sardinian Museum of Anthropology and Ethnography of the University of Cagliari.

**Seulo 68 Neo-Iron_47 (*Sardinia_Nuragic_BA_contam*, I3664).** Fragment of bone diaphisis, C14 dated to 1880-1700 calBCE (3470±20 BP, PSUAMS-2390).

**SEULO 59 (*Sardinia_Nuragic_BA*, I3743).** Tooth, C14 dated to 2134-1947 calBCE (3655±25 BP, PSUAMS-2076).

**Seulo_Ind.B (*Sardinia_Nuragic_BA10365*, I10365).** Petrous bone, C14 dated to 1643-1263 calBCE (3190±80 BP, Beta-37705).

*Usellus, Sardinia, Italy ( ⊕ 39.81, 8.84)*

Motrox 'e Bois (Usellus) is a collective burial monument typical of the so-called "giants' tombs" of Nuragic Sardinia. It is located in the historical-geographical region of Arborea, central western Sardinia, at 289 m above sea level. The excavations took place in 1957. Based on the study of mandibles and isolated teeth, a minimum number of 42 individuals was estimated.

The skeletal material has been dated back to the Final Bronze Age by radiocarbon analysis. The osteological collection of Motrox 'e Bois burial is housed in the Sardinian Museum of Anthropology and Ethnography of the University of Cagliari[16].

**Motrox'e Bois-3 (*Sardinia_Nuragic_BA_lowcov*, I10368).** Petrous bone, C14 dated to 1150-800 BCE [layer date of 1126-825 calBCE (2810±60 BP, Beta-82329)].

**MSAE 6927 (Motrox'e Bois) (*Sardinia_Nuragic_BA_lowcov*, I10367).** Petrous bone, Context dated to 1150-800 BCE based on a C14 date of 1126-825 calBCE (2810±60 BP, Beta-82329) from the same layer.

**MSAE 8784 (Motrox 'e Bois) (*Sardinia_IA*, I10366). Petrous bone, C14 dated to 391-209 calBCE (2250±20 BP, PSUAMS-4874)**

*Grotta Colombi, Sardinia, Italy ( ⊕ 39.182, 9.163)*

Sant'Elia Cape (Cagliari, Southern Sicily) harbors extensive archaeological material in the caves of its Miocene limestone cliff. It includes the well-preserved Cala Mosca stratigraphic section, which represents the type locality of the so-called Tyrrhenian plain[17]. At Marine Isotope State (MIS) 5.5, high sea levels caused the Sant'Elia hill to be an island separated from the Cagliari coast by about 1.5 km of sea. During successive glacial periods when sea levels were lower, the marine shelf surrounding the Sant'Elia promontory was largely exposed. For the last 7000-5000 years, the coastline has followed a similar course to the present one.

The Grotta dei Colombi (Pigeon's Cave) is one of the five natural caves located on the slope of the promontory, deriving its name from the many wild pigeons that have nested there in the past. This large cave has been known since ancient times, and opens today just above sea level.

The cave entrance, triangular in shape, can only be accessed from the sea. The internal hall, approximately circular in shape, is about 30 meters wide and 20 meters high.

During the Late Neolithic (if not earlier) the cave has been either temporally inhabited or visited for funeral use by people reaching it by boats as documented by the presence of a human skeleton and fragments of pre-Nuragic ceramic pottery (Ozieri Culture, 3200-2800 BCE) in the lowermost levels of the sedimentary sequence filling the cave. The uppermost levels contain a variety of Nuragic pottery dated to the 8th century BCE[18-21].

The cave was still occasionally visited by fishermen and hunters even in recent times.

The human remains analyzed here were collected during a geological survey by the Italian researcher A. Malatest at different points on the surface floor of the Grotta dei Colombi cave together with some fragments of ceramic pottery and few fragments of bird (mainly *Columba livia*) and mammal bones (*Capra hircus, Ovis aries*, medium-sized deer).

**GC3 (*Sardinia_LateAntiquity*, I12220).** Long bone, C14 dated to 566-640 calCE (1465±15 BP, PSUAMS-5283).

**GC4 (*Sardinia_LateAntiquity12221*, I12221).** Bone, context dated to 200-700 BCE based on C14 dates from other individuals at the same site.

**GC5 (*Sardinia_LateAntiquity*, I12222).** Bone, context dated to 200-700 BCE based on C14 dates from other individuals at the same site.

**GC1 (*Sardinia_LateAntiquity*, I12223).** Tooth, 14 dated to 256-403 calCE (1700±25 BP, PSUAMS-5427).

*Contrada Paolina, Castellucciana, Sicily, Italy ( ⊕ 36.883, 14.571)*

The skeletal remains come from the district of Paolina, in the hinterland of Camarina in Ragusa Province, Southern Sicily, Italy. The tomb complex, consisting of three tombs, is located on a low hill between the last slope of the Ragusa plateau down towards the sea. The excavation was conducted between April and May 1977 by the Archaeological Superintendence of Syracuse. Three artificial tombs are present, excavated in slightly coherent limestone. The human remains come from Tomb 2 with a diameter of 2.40 and height of 1.32, a “grotticella” tomb of the "oven type” in the common typology of the Castelluccian tombs[22]. In previous studies, the presence of at least 76 individuals has been estimated, including adults, and infants of both sexes (determined anthropologically)[23-27]. The multiple depositions were grouped along the tomb chamber walls, interpreted as subsequent burials in different times probably by family or clan (common rite in coeval sites[22]) and a single primary undisturbed burial in the middle of the room.

**PP B (*Sicily_EBA_contam*, I7743).** Context dated to 3000-1600 BCE.

**PP E (*Sicily_EBA_lowcov*, I7772).** Context dated to 3000-1600 BCE.

**PP F (*Sicily_EBA_lowcov*, I7773).** Context dated to 3000-1600 BCE.

**PP G (*Sicily_EBA_contam*, I7774).** Context dated to 3000-1600 BCE.

**PP H (*Sicily_EBA_lowcov*, I7775).** Context dated to 3000-1600 BCE.

**PP I (*Sicily_EBA_lowcov*, I7776).** Context dated to 3000-1600 BCE.

**PP IX (*Sicily_EBA_contam*, I7735).** Context dated to 3000-1600 BCE.

**PP K (*Sicily_EBA*, I7796).** Context dated to 3000-1600 BCE.

**PP O (*Sicily_EBA*, I7800).** Context dated to 3000-1600 BCE.

**PP V (*Sicily_EBA, I7807*).** Context dated to 3000-1600 BCE.

**PP XIII (*Sicily_EBA*, I7738).** Context dated to 3000-1600 BCE.

*Isnello, Sicily, Italy ( ⊕ 37.95, 14.083)*

The Cave Abisso del Vento, of karstic origin, is located in the locality "Ficuzedde" in the Municipality of Isnello (PA), and it is one of the most complex cave systems in Sicily in the mountain districts of Pizzo Dipilo (1385 m asl) and Pizzo Carbonara (m 1979 above sea level) in the eastern Madonie area. The cave has produced anthropological materials since the nineteenth century, classifiable as Eneolithic-Early Bronze Age and currently kept at the Minà Palumbo Museum in Castelbuono. The sample we analyzed derives from a multiple burial just below the entrance of the cave. The anthropological analysis sex determination agreed with the genetic determination.

**Abisso del Vento 2017 (Grotto) (*Sicily_EBA8561,* I8561).** C14 dated to 2346-2199 calBCE (3825±20 BP, PSUAMS-4873).

*Vallone Inferno, Sicily, Italy (⊕ 37.872, 13.935)*

Vallone Inferno rock-shelter is an archaeological site located in the Madonie mountain range in Sicily[28]. Archaeological excavation has provided a long prehistoric and historic sequence from the Neolithic to the Medieval period. Of the four stratigraphic complexes, complex 3 has provided almost all the archaeological remains. 14C AMS dates, obtained from five samples, place the human activities between the Middle Neolithic, radiocarbon dated to ~5460 calBCE and the Medieval period ~644 calCE[29]. A persistent use of the shelter for pastoral (herding) activities has been shown from macrofaunal and plant remains. The aridification and the opening of the landscape from the base to the top of the sequence has been observed by the analysis of environmental data, obtained from microvertebrate and archaeobotanical remains.

Within the archaeological layers of the Early Bronze Age, the main cultural attribution of the ceramic is to the Castelluccian culture, with the sporadic presence (one ceramic fragment) of Bell Beaker associated pottery.

Several human remains were found within the Early Bronze Age layers, with a preponderance of dental remains (76%)[28].

***VALINF (Sicily_BA_lowcov,* I4383).** Long bone, context dated to 2600-2300 BCE based on its relationship to a radiocarbon date on bone in layer 3.4.g of 2601-2309 calBCE (3948±35 BP, DSH-1976), and a radiocarbon date on seed in layer 3.4.b of 1643-1411 calBCE (3244±42 BP, DSH-2815).

**2 - *qpWave* clade analysis**

We used *qpWave*[30] from ADMIXTOOLS[31] to cluster individuals. We used a p>0.01 threshold as a criterion for clustering, *allsnps:YES*, and the following “Right” populations:

*Mbuti.DG, Ust_Ishim, CHG, EHG, ElMiron, Vestonice16, MA1, Israel_Natufian, Jordan_PPNB, Anatolia_Neolithic, WHG, Iran_Ganj_Dareh_Neolithic, Yamnaya_Samara*

The results of this analysis are presented in **Fig. 3** and summarized below. Pairs of individuals that passed this *qpWave* analysis at the p>0.01 threshold and were from the same region and chronological period were lumped for analysis with a few exceptions.

After identifying all pairs of chronologically/geographically grouped individuals that gave evidence of being different in ancestry at the p<0.01 level, we further tested them using the same method for a difference in ancestry compared to other individuals in the grouping.

<u>Bronze Age Balearic Islands:</u>

Only one pair of individuals was detected as genetically inhomogeneous by *qpWave*: *Mallorca_EBA-Menorca_LBA* (p=0.002). (**Supplementary Table 1**). While *Formentera_MBA* was consistent with being a clade with each of the other two, two $f_4$-statistic symmetry tests quoted in the main text reveal that *Menorca_LBA* shared alleles at different rates with both *Iberia_Chalcolithic* and *Sardinia_Nuragic_BA*. Based on this and also the chronological differences among the three Balearic island samples, we analyzed them individually.

**Supplementary Table 1:** P-values for the *qpWave* models between the Balearic individuals. (In parentheses we give P-values based on using a more sensitive outgroup set adding in more closely related groups; that is, we add in *Sardinia_Nuragic_BA* and *Sicily_EBA*.)

| | Mallorca_EBA | Formentera_MBA | Menorca_LBA |
|---|---|---|---|
| **Mallorca_EBA** | | 0.680 (0.685) | 0.002 (0.001) |
| **Formentera_MBA** | 0.680 (0.685) | | 0.748 (0.453) |
| **Menorca_LBA** | 0.002 (0.001) | 0.748 (0.453) | |

The Iron Age Phoenician individual from Ibiza from [32] did not form a clade with any of the three preceding individual from our study (*Mallorca_EBA* p=$4.0\times10^{-12}$, *Formentera_MBA* p=$1.4\times10^{-5}$, *Menorca_LBA* p<$10^{-12}$), but instead did so with the Late Antiquity Sicilian individuals (p-value always above 0.154) that possessed large amounts of Iranian-related ancestry (**Fig. 3**), clearly showing not only a foreign origin but also the arrival of ancestries not previously seen in the Balearic Islands.

When we looked at modern Balearic individuals they only formed a clade with *Sardinia_LateAntiquity12221* (p=0.102) and *Sicily_EBA11443* (p=0.018) (**Fig. 3**).

<u>Nuragic Bronze Age Sardinians:</u>

Individual I10365 was not consistent with forming a clade with 3 of the other Nuragic Sardinian individuals from our dataset, and the model I3741-I10554 also produced a p-value below the threshold of 0.010 (**Supplementary Table 2**).

**Supplementary Table 2:** P-values for *qpWave* between the Nuragic Sardinian individuals. (In parentheses we give P-values based on using a more sensitive outgroup set adding in more closely related groups; that is, we add *Sicily_EBA, Bell_Beaker_Iberia, Bell_Beaker_Iberia_highsteppe, France_Bell_Beaker, France_Bell_Beaker_lowsteppe,* and *Anatolia_EBA.*)

| | I10365 | I10364 | I10552 | I10553 | I10554 | I3642 | I3741 | I3743 |
|---|---|---|---|---|---|---|---|---|
| **I10365** | | 0.044 (0.084) | 4.23E-05 (9.11E-05) | 1.64E-04 (9.98E-05) | 0.070 (7.89E-03) | 0.077 (0.137) | 0.019 (0.025) | 6.55E-03 (0.021) |
| **I10364** | 0.044 (0.084) | | 0.155 (0.263) | 0.171 (0.045) | 0.491 (0.173) | 0.619 (0.528) | 0.033 (0.045) | 0.362 (0.459) |
| **I10552** | 4.23E-05 (9.11E-05) | 0.155 (0.263) | | 0.027 (0.026) | 0.050 (0.073) | 0.290 (0.358) | 0.791 (0.123) | 0.179 (0.337) |
| **I10553** | 1.64E-04 (9.98E-05) | 0.171 (0.045) | 0.027 (0.026) | | 0.096 (0.028) | 0.107 (0.121) | 0.100 (3.37E-03) | 0.151 (0.335) |
| **I10554** | 0.070 (7.89E-03) | 0.491 (0.173) | 0.050 (0.073) | 0.096 (0.028) | | 0.181 (0.106) | 9.82E-03 (1.50E-04) | 0.046 (0.019) |
| **I3642** | 0.077 (0.137) | 0.619 (0.528) | 0.290 (0.358) | 0.107 (0.121) | 0.181 (0.106) | | 0.638 (0.151) | 0.711 (0.857) |
| **I3741** | 0.019 (0.025) | 0.033 (0.045) | 0.791 (0.123) | 0.100 (3.37E-03) | 9.82E-03 (1.50E-04) | 0.638 (0.151) | | 0.101 (0.015) |
| **I3743** | 6.55E-03 (0.021) | 0.362 (0.459) | 0.179 (0.337) | 0.151 (0.335) | 0.046 (0.019) | 0.711 (0.857) | 0.101 (0.015) | |

We further tested the three individuals that were involved in these p<0.01 outcomes against the pool of Nuragic individuals that were all mutually consistent with forming a clade. Only individual I10365 is not consistent with forming a clade with the grouped individuals at p<0.01. We therefore treated only this individual as an outlier in subsequent analysis (**Supplementary Table 3**).

**Supplementary Table 3:** P-values for the *qpWave* models between the Nuragic Sardinian individuals with some evidence of ancestry heterogeneity, against all other individuals.

| | I3642+I3743+I10364+I10552+I10553 |
|---|---|
| **I10365** | 2.54E-05 |
| **I10554** | 0.043 |
| **I3741** | 0.116 |

Iron Age Sardinian (I10366):

The Iron Age individual from Sardinia (391-209 calBCE) was not consistent with forming a clade with any of the individuals from the Nuragic Bronze Age, but was consistent with forming a clade with two of the four individuals from Late Antiquity (I12220, p=0.109; I12222, p=0.114). This suggests arrival in Sardinia of new ancestry types at least by the Iron Age, potentially coinciding with the period of Phoenician settlement. We analyzed I10366 separately.

Late Antiquity Sardinians:

All four individuals from this period were consistent with forming a clade based on *qpWave* (**Supplementary Table 4**), despite the fact that in PCA, I12221 is an outlier from the cluster of the other 3 individuals (**Fig. 2**). Based on the PCA and the different performance of I12221 in *qpAdm* modeling (below), we made an exception to the strict *qpWave* sample grouping procedure and analyzed the individuals of this period as a main grouping of three individuals *Sardinia_LateAntiquity* along with an outlier individual *Sardinia_LateAntiquity12221*.

**Supplementary Table 4:** P-values for the *qpWave* models between the Late Antiquity Sardinians. (In parentheses we give P-values based on using a more sensitive outgroup set adding in more closely related groups; that is, we add *Sardinia_Nuragic_BA, Sardinia_Nuragic_BA10365,* and *Sardinia_IA*.)

| | **I12220** | **I12222** | **I12223** | **I12221** |
|---|---|---|---|---|
| **I12220** | | 0.638 (0.417) | 0.827 (0.797) | 0.587 (0.178) |
| **I12222** | 0.638 (0.417) | | 0.352 (0.440) | 0.274 (0.056) |
| **I12223** | 0.827 (0.797) | 0.352 (0.440) | | 0.959 (0.816) |
| **I12221** | 0.587 (0.178) | 0.274 (0.056) | 0.959 (0.816) | |

We finally analyzed modern Sardinians (*Sardinian*) and found that they were not consistent with forming a clade with the main cluster of Nuragic Bronze Age individuals (*Sardinia_Nuragic_BA*) (p<10-12). However, they were consistent with forming clades with the outlier *Sardinia_Nuragic_BA10365* (p=0.133). Thus, modern Sardinian individuals cannot have been direct descendants without admixture of the main population of which most of the Nuragic individuals analyzed in our study were a part.

Early Bronze Age Sicily:

Among the Sicilian Early Bronze assemblage individuals, I11443 from Buffa Cave (*Sicily_EBA11443*) and individual I8561 from Isnello (*Sicily_EBA8561*) were not consistent with forming a clade with the remaining Early Bronze Age Sicilians in the great majority of tests. We therefore treat them as separate outliers (**Supplementary Table 5**).

**Supplementary Table 5:** P-values for the *qpWave* models between Early Bronze Age Sicilians. (In parentheses we give P-values based on using a more sensitive outgroup set adding in more closely related groups; that is, we add *Sardinia_Nuragic_BA, Bell_Beaker_Iberia_highsteppe, France_Bell_Beaker, France_Bell_Beaker_lowstepp,* and *Anatolia_EBA.*)

| | I11442 | I3122 | I3123 | I7796 | I7800 | I7807 | I11443 | I8561 |
|---|---|---|---|---|---|---|---|---|
| **I11442** | | 0.298 (0.154) | 0.110 (0.114) | 0.922 (0.655) | 0.126 (0.288) | 0.022 (0.051) | 5.31E-31 (2.88E-29) | 1.10E-10 (5.72E-10) |
| **I3122** | 0.298 (0.154) | | 0.015 (0.008) | 0.812 (0.803) | 0.364 (0.410) | 0.594 (0.286) | 7.92E-40 (5.08E-41) | 3.57E-11 (6.11E-13) |
| **I3123** | 0.110 (0.114) | 0.015 (0.008) | | 0.054 (0.087) | 0.760 (0.901) | 0.004 (0.015) | 2.95E-15 (4.81E-15) | 0.006 (0.001) |
| **I7796** | 0.922 (0.655) | 0.812 (0.803) | 0.054 (0.087) | | 0.386 (0.384) | 0.926 (0.628) | 3.34E-23 (3.28E-21) | 4.12E-07 (5.05E-08) |
| **I7800** | 0.126 (0.288) | 0.364 (0.410) | 0.760 (0.901) | 0.386 (0.384) | | 0.471 (0.593) | 6.56E-07 (5.19E-06) | 0.017 (0.021) |
| **I7807** | 0.022 (0.051) | 0.594 (0.286) | 0.004 (0.015) | 0.926 (0.628) | 0.471 (0.593) | | 1.32E-24 (2.33E-24) | 7.76E-07 (1.75E-06) |
| **I11443** | 5.31E-31 (2.88E-29) | 7.92E-40 (5.08E-41) | 2.95E-15 (4.81E-15) | 3.34E-23 (3.28E-21) | 6.56E-07 (5.19E-06) | 1.32E-24 (2.33E-24) | | 1.41E-07 (1.95E-07) |
| **I8561** | 1.10E-10 (5.72E-10) | 3.57E-11 (6.11E-13) | 0.006 (0.001) | 4.12E-07 (5.05E-08) | 0.017 (0.021) | 7.76E-07 (1.75E-06) | 1.41E-07 (1.95E-07) | |

The pair I3123-I7807 produced a p<0.01 so we compared both individuals against the group that gave no evidence of outliers (**Supplementary Table 6**). Individual I3123 was not consistent with forming a clade with the main grouping (p=0.004) so we treated him as an outlier (*Sicily_EBA3123*). We grouped the remaining five Sicilian Early Bronze Age individuals as *Sicily_EBA* for our main analyses.

**Supplementary Table 6:** P-values for the *qpWave* models between Early Bronze Age Sicilian individuals with some evidence of ancestry heterogeneity, against all other individuals.

| | I3122+I7796+I7800+I11442 |
|---|---|
| **3123** | 0.004 |
| **11443** | 4.13E-46 |
| **I8561** | 1.67E-16 |
| **I7807** | 0.245 |

Middle Bronze Age Sicily:

The pair of individuals I3124-I4109 did not form a clade (p<0.010) (**Supplementary Table 7**). Based on this and heterogeneity in *qpAdm* results for all three Middle Bronze Age individuals we went beyond the *qpWave* inference and analyzed all three separately.

**Supplementary Table 7:** P-values for *qpWave* models between Middle Bronze Age Sicilians. (In parentheses we give P-values based on using a more sensitive outgroup set adding in more closely related groups; that is, we add *Sicily_EBA, Sicily_EBA3123, Sicily_EBA8561* and *Sicily_EBA11443.*)

| | I3124 | I3125 | I4109 |
|---|---|---|---|
| **I3124** | | 0.044 (0.072) | 4.77E-04 (0.003) |
| **I3125** | 0.044 (0.072) | | 0.234 (0.356) |
| **I4109** | 4.77E-04 (0.003) | 0.234 (0.356) | |

Late Bronze Age Sicily:

All Late Bronze Age individuals were consistent with forming a clade with each other (**Supplementary Table 8**), and so we pooled them as *Sicily_LBA*.

**Supplementary Table 8:** P-values for *qpWave* models between Late Bronze Age Sicilians. (In parentheses we give P-values based on using a more sensitive outgroup set adding in more closely related groups; that is, we add *Sicily_EBA, Sicily_EBA3123, Sicily_EBA8561* and *Sicily_EBA11443, Sicily_MBA3124, Sicily_EBA3125* and *Sicily_EBA4109.*)

| | I10371 | I10372 | I10373 | I3876 | I3878 |
|---|---|---|---|---|---|
| **I10371** | | 0.026 (0.064) | 0.027 (0.113) | 0.193 (0.232) | 0.034 (0.066) |
| **I10372** | 0.026 (0.064) | | 0.091 (0.038) | 0.022 (0.009) | 0.045 (0.022) |
| **I10373** | 0.027 (0.113) | 0.091 (0.038) | | 0.443 (0.706) | 0.957 (0.685) |
| **I3876** | 0.193 (0.232) | 0.022 (0.009) | 0.443 (0.706) | | 0.647 (0.275) |
| **I3878** | 0.034 (0.066) | 0.045 (0.022) | 0.957 (0.685) | 0.647 (0.275) | |

We found that all modern Sicilians were consistent with forming a clade with *Ibiza_Phoenician* (p=0.013), although marginally, as well as with all *Sardinia_LateAntiquity* individuals (*Sardinia_LateAntiquity12220*, p=0.041, *Sardinia_LateAntiquity12221*, p=0.097, *Sardinia_LateAntiquity12222*, p=0.056, *Sardinia_LateAntiquity12220*, p=0.240).

##### 3 - *qpAdm* modelling

We used *qpAdm* (https://github.com/DReichLab)[30] to infer proportions of ancestry in various test populations relative using as surrogates both very distantly related populations ("distal modeling") and closely related populations ("proximal modeling").

*qpAdm* works by computing all possible statistics of the form $f_4(Left_i, Left_j; Right_k, Right_l)$ relating a *Test* population and a set of groups proposed to be clades with the sources of ancestry in the *Test* population (together, these constitute the "Left" populations in *qpAdm*), and a set of outgroups chosen to be differentially related to the *Left* populations ("Right" populations). The method produces a p-value for the fit of the model, as well as admixture proportions[30,33].

We used the following 9 populations/individuals as our based "Right" set:

*Mbuti.DG, Ust_Ishim, CHG, EHG, ElMiron, Vestonice16, MA1, Israel_Natufian, Jordan_PPNB*

For distal modeling, following the strategy of [34], we began by investigating the ancestry of each Test individual/population as derived from a set of four geographical and chronologically distal populations in order to identify and quantify the influence of four major ancestries known to have contributed to Bronze Age Europeans. We used a pool of *Anatolia_Neolithic* individuals to represent the ancestry brought into Europe by western Anatolian farmers around 8500 years ago. We represented *WHG* (Western European hunter-gatherers) by 5 individuals from Spain (*Chan.SG* and *LaBrana1.SG*), Hungary (I1507 and I4971), and Luxembourg (*Loschbour.DG*). We used *Yamnaya_Samara* individuals as a proxy for the ancestry brought into Europe by Steppe pastoralist groups after around 3000 BCE. We used *Iran_Ganj_Dareh_Neolithic* as a proxy for the Western Zagros farmer-related ancestry (Iranian-related ancestry) that has been documented to have contributed to Middle to Late Bronze Age Aegean populations by at least ~2000 BCE.

In our distal modeling, we used *Anatolia_Neolithic* and *WHG* as base ancestries for European populations. If a parsimonious model of mixture of these two groups did not fit at the p>0.05 level, we then tried adding in either *Yamnaya_Samara* or *Iran_Ganj_Dareh_Neolithic* as a 3-way model. When both models fit we used a "model competition" approach, moving the other potential source population into the Right set of populations. In practice, we found that this always resulted in a single fitting parsimonious model for each analysis grouping. Finally, if neither of these 3-way mixture models worked, we tried fitting a 4-way model adding in both of these groups.

##### 3.1 - Results of Distal Modeling

We ran *qpAdm* with the groupings obtained as described in the previous section (**Supplementary Table 9**).

In the Balearic islands, we observe a decrease in *Yamnaya_Samara*-related ancestry from the Early to the Late Bronze Age (36.9 ± 4.2% to 23.1 ± 3.6%), along with a concomitant increase in

*Anatolia_Neolithic*-related ancestry. We do not need to model any Iranian-related ancestry to fit the Balearic individuals.

We tried to model the Phoenician individual from [32] using the same 4 distal sources but no models produced valid results. When we added *Morocco_LN* as a fifth possible source, however, we obtained a good two-way fit for a model with 18.8 ± 7.9% *Anatolia_Neolithic* and 81.2 ± 7.9% *Morocco_LN* ancestry (p=0.141).

Modern Balearic island individuals do not quite fit a 4-way model (p=0.037), and the addition of *Morocco_LN* to the "Right set" clearly breaks the model ($p=8.7x10^{-9}$) suggesting North African ancestry as in the Phoenician individual. By adding *Morocco_LN* as a source we found that the only working model at the p>0.05 level (p=0.139) involved 5-way mixture with 25.4 ± 6.2% *Anatolia_Neolithic*, 8.2 ± 2.2% *WHG*, 13.6 ± 6.1% *Iran_Ganj_Dareh_Neolithic*, 26.3 ± 2.2% *Yamnaya_Samara* and 26.6 ± 12.3% *Morocco_LN*.

In Nuragic Sardinia, both the grouped individuals and the outlier are consistent with simple mixtures of *Anatolia_Neolithic* and *WHG* ancestries (the estimated proportion of *Anatolia_Neolithic* is 82.5 ± 1.1% for *Sardinia_Nuragic_BA*, and 82.5 ± 1.1% for *Sardinia_Nuragic_BA10365*). We also find that modern Sardinians harbor ancestry that is more closely related to *Sardinia_Nuragic_BA* than to *Anatolia_Neolithic,* as when we add modern Sardinians (*Sardinian*) to the outgroup set in our distal *qpAdm* modeling, we reject the model at high significance ($p<10^{-12}$). This is consistent with some degree of continuity between Bronze Age and modern Sardinians.

The Iron Age individual can only fit if we also model in Iranian-related ancestry (p=0.167; the fits is p=0.0066 for the 2-way model that works for Nuragic individuals, and p=0.037 for the 3-way model that uses *Yamnaya_Samara* instead of *Iran_Ganj_Dareh_Neolithic* as a source (**Supplementary Table 9**). This evidence of Iranian-related ancestry is the earliest genome-wide ancient DNA evidence for eastern ancestry in Sardinia. The radiocarbon date of 391-209 calBCE for this individual places it after the date of Phoenician influence in Sardinia, making Phoenicians a plausible source for this ancestry, a scenario that is also supported by previously reported mitochondrial ancient DNA evidence documenting Phoenician-associated mitochondrial DNA haplogroups in Sardinia[35]. If the ancestry source was Phoenician, it is possible that this individual harbored North African admixture but when we added Late Neolithic Moroccans (*Morocco_LN*)[36] to the "Right" set of outgroups, the model still passed p>0.05 and hence we cannot establish a contribution of North Africa to the limits of our resolution (p=0.236) (**Supplementary Table 9**). As in the case of the Nuragic Bronze Age Sardinians, the addition of modern Sardinians (*Sardinian*) to the "Right" outgroup set also makes the distantl model fail ($p<10^{-12}$), supporting the hypothesis that *Sardinia_IA* harbors ancestry from lineages that also contributed to modern Sardinians.

For the Sardinian Late Antiquity individuals (~200-700 CE) we find no working 3-way models at a significance of p>0.05 for the pool of all four individuals, so we tried removing individual I12221, who in PCA and ADMIXTURE has evidence of ancestry different from that of the other three individuals (**Fig. 2**). Without I12221, the model with *Anatolia_Neolithic* (65.4 ± 5.3%), *WHG* (5.0 ± 2.6%) and *Iran_Ganj_Dareh_Neolithic* (29.6 ± 4.6%) yields a plausible p-value of 0.104, thus confirming the presence of Iranian-related ancestry in Sardinia already documented in the Iron Age individual. In contrast, the outlier I12221 (*Sardinia_LateAntiquity12221*) fits with an alternative model of *Anatolia_Neolithic* (66.7 ± 5.5%) and *Yamnaya_Samara* (33.3 ± 5.5%) ancestry (the three-way fit including WHG yielded a negative estimate of -11.9 ± 5.4%, so we exceptionally modeled this individual without *WHG* and found that the model of *Anatolia_Neolithic* (66.7 ± 5.5%) and *Yamnaya_Samara* admixture worked (33.3 ± 5.5%) (p=0.067) **(Supplementary Table 9**). These results provide further justification for analyzing I12221 separately. The addition of *Morocco_LN* to the "Right" did not cause model failures for *Sardinia_LateAntiquity* (p=0.148) and *Sardinia_LateAntiquity12221* (p=0.097), showing that as with the Sardinian Iron Age individual, there is no significant evidence of North African ancestry to these individuals.

Our dataset of 4 modern Sardinians (*Sardinian.DG*[31]) can only be modeled with additional *Iran_Ganj_Dareh_Neolithic*-related ancestry (the estimated proportion of *Anatolia_Neolithic* is 58.1 ± 2.4% and the estimated proportion of *Iran_Ganj_Dareh_Neolithic* is 28.6% ± 2.2%). This is a surprising result in light of the literature, which has not previously inferred Iranian-related ancestry in present-day Sardinians, and hence we repeated the analysis using a larger set of 27 Sardinians genotyped on the Affymetrix Human Origins SNP array (where we have data at only about half the SNPs but where the larger sample size compensates in statistical power) (Sardinian)33. In this analysis, no 3-way mixture model fits but a 4-way mixture model fits which again includes Iranian-related ancestry: the estimated proportion of *Anatolia_Neolithic* is 61.4% ± 1.6%, the estimated proportion of *Iran_Ganj_Dareh_Neolithic* is 19.1% ± 1.9%, and the estimated proportion of *Yamnaya_Samara* is 10.0% ± 1.6% (**Supplementary Table 10**). Despite the significant p-values, we found that by adding *Morocco_LN* to the "Right" all these models failed, suggesting a North African influence in present-day Sardinians that is not detected in any of the ancient individuals in our dataset. The 5-way model with *Morocco_LN* (16.1 ± 8.4%) as a source produces a valid model (p=0.235). This finding may be related to the observation of North African admixture in the ancestry of present-day Sardinians by Hellenthal and colleagues which was inferred to have an average admixture date of ~630 CE[37], as well as findings of sub-Saharan African admixture which could have been mediated by North African admixture into Sardinia[38-40].

In Sicily, the Middle Neolithic individuals can be modeled as 2-way mixtures of *Anatolia_Neolithic* (88.9 ± 1.2%) and *WHG*. We can fit a similar model without any requirement for eastern ancestry in data from a Bell Beaker-associated individual for whom very low coverage data were originally reported in [41] and whom we re-analyze here with increased quality data. By the Early Bronze Age, however, we definitively detect *Yamnaya_Samara* related ancestry in *Sicily_EBA* (9.1 ± 2.2%) and the outliers *Sicily_EBA11443* (40.2 ± 3.5%), *Sicily_EBA8561* (23.3 ± 3.5%), and *Sicily_EBA3123* (14.1 ± 3.4%).

Two Middle Bronze Age Sicilian individuals can only be fit parsimoniously with models requiring Iranian-related ancestry: *Sicily_MBA3125* (18.0 ± 3.6%, p=0.33) and *Sicily_MBA4109* (14.9 ± 3.9%, p=0.24) while models that use only *Yamnaya_Samara* ancestry as a source do not fit (p=0.032 and p=0.037 respectively, which are below our threshold of $p<0.05$). The other Middle Bronze Age Sicilian, *Sicily_MBA3124*, initially produced valid 3-way models with either *Yamnaya_Samara* or *Iran_Ganj_Dareh_Neolithic* but when we repeated the tests adding the other population to the "Right" outgroup set then only the model with *Yamnaya_Samara* produced a fit (13.3 ± 3.3%, p=0.52) (the model with *Iran_Ganj_Dareh_Neolithic* was rejected at 0.002). These results are consistent with *qpWave* in suggesting ancestry heterogeneity in the Middle Bronze Age Sicilians. In light of the ubiquitous presence of Steppe ancestry in the Early Bronze Age and its detection in one of the Middle Bronze Age individuals, we hypothesize that with more statistical power we might detect 4-way mixture in the Middle Bronze Age with Iranian-related ancestry.

In Late Bronze Age Sicily at the site of Marcita, our *qpAdm* models required only the presence of *Anatolia_Neolithic* (80.2 ± 1.8%), *WHG* (5.3 ± 1.6%), and *Yamnaya_Samara* (14.5 ± 2.2%), with no requirement of Iranian-related ancestry. Thus, the Iranian-related ancestry detected in earlier periods was proportionally less important in Late Bronze Age Marcita.

As with the Balearic individuals we could not model modern Sicilians using a threshold of $p>0.05$, or even with a more permissive $p>0.01$ threshold. There is clearly also a North African influence, however, as we identify a working model for the 4-way model with 24.8 ± 4.3% *Anatolia_Neolithic*, 12.1 ± 3.1% *Iran_Ganj_Dareh_Neolithic*, 19.8 ± 1.4% *Yamnaya_Samara*, and 43.3 ± 6.1% *Morocco_LN* (p=0.334).

Adding *Morocco_LN* to the "Right" did not break any of the Sicilian models, suggesting low levels of such ancestry in the analyzed ancient individuals (**Supplementary Table 9**).

**Supplementary Table 9:** Admixture proportions for the most parsimonious *qpAdm* models of the populations/individuals from this study fit as derived from groups related to four sources - *Anatolia_Neolithic* (1), *WHG* (2), *Iran_Ganj_Dareh_Neolithic* (3), and Y*amnaya_Samara* (4). In the final column, we show the p-value of the most parsimonious model (shaded in green) when we add *Morocco_LN* to the "Right" outgroup set. "Right" outgroup set: *Mbuti.DG, Ust_Ishim, CHG, EHG, ElMiron, Vestonice16, MA1, Israel_Natufian, Jordan_PPNB*. These data are used for **Fig. 4b**.

| *Test* | P-value | Admixture Sources and Proportions | | | | Standard Error | | | | P-values for key models | | | P-value of the most parsimonious model after adding Morocco_LN to the "Right" |
|---|---|---|---|---|---|---|---|---|---|---|---|---|---|
| | | 1 | 2 | 3 | 4 | SE1 | SE2 | SE3 | SE4 | 1+2 | 1+2+3 | 1+2+4 | |
| *Mallorca_EBA* | 0.665 | 0.454 | 0.177 | - | 0.369 | 0.036 | 0.033 | - | 0.042 | 5.54E-14 | 1.98E-09 | 0.665 | 0.758 |
| *Formentera_MBA* | 0.550 | 0.597 | 0.140 | - | 0.263 | 0.044 | 0.038 | - | 0.051 | 5.80E-05 | 0.002 | 0.550 | 0.629 |
| *Menorca_LBA* | 0.299 | 0.594 | 0.174 | - | 0.231 | 0.028 | 0.027 | - | 0.036 | 4.01E-08 | 3.45E-06 | 0.299 | 0.370 |
| *Ibiza_Phoenician* | 3.50E-04 | 0.631 | 0.032 | 0.337 | - | 0.057 | 0.029 | 0.051 | - | 1.10E-12 | 3.50E-04 | 9.52E-02 | 5.66E-04 |
| *Balearic (modern)* | 0.037 | 0.373 | 0.118 | 0.258 | 0.251 | 0.026 | 0.016 | 0.031 | 0.026 | 2.51E-120 | 1.79E-14 | 1.26E-18 | 8.73E-09 |
| *Sardinia_Nuragic_BA* | 0.265 | 0.825 | 0.175 | - | - | 0.011 | 0.011 | - | - | 0.265 | 0.235 | 0.873 | 0.058 |
| *Sardinia_Nuragic_BA10365* | 0.064 | 0.854 | 0.146 | - | - | 0.022 | 0.022 | - | - | 0.064 | 0.389 | 0.436 | 0.099 |
| *Sardinia_IA* | 0.168 | 0.759 | 0.122 | 0.119 | - | 0.040 | 0.022 | 0.037 | - | 0.007 | 0.168 | 0.037 | 0.236 |
| *Sardinia_LateAntiquity* | 0.104 | 0.654 | 0.050 | 0.296 | - | 0.053 | 0.026 | 0.046 | - | 9.47E-09 | 0.104 | 2.00E-06 | 0.148 |
| *Sardinia_LateAntiquity12221* | 0.067 | 0.667 | - | - | 0.333 | 0.055 | - | - | 0.055 | 3.71E-07 | 0.001 | 0.243 | 0.097 |
| *Sardinian.DG (modern)* | 0.163 | 0.581 | 0.134 | 0.286 | - | 0.024 | 0.013 | 0.022 | - | 2.45E-40 | 0.163 | 6.43E-15 | 7.40E-05 |
| *Sardinian* (modern)* | 0.116 | 0.614 | 0.095 | 0.191 | 0.100 | 0.016 | 0.010 | 0.019 | 0.016 | 1.82E-92 | 4.88E-07 | 7.07E-26 | 8.96E-13 |
| *Sicily_MN* | 0.151 | 0.889 | 0.111 | - | - | 0.012 | 0.012 | - | - | 0.151 | 0.105 | 0.230 | 0.188 |
| *Sicily_Bell_Beaker* | 0.474 | 0.831 | 0.169 | - | - | 0.079 | 0.079 | - | - | 0.474 | 0.374 | 0.265 | 0.395 |
| *Sicily_EBA* | 0.265 | 0.838 | 0.071 | - | 0.091 | 0.019 | 0.017 | - | 0.022 | 0.001 | 0.016 | 0.265 | 0.132 |
| *Sicily_EBA3123* | 0.623 | 0.752 | 0.107 | - | 0.141 | 0.029 | 0.025 | - | 0.034 | 0.003 | 0.015 | 0.623 | 0.156 |
| *Sicily_EBA11443* | 0.824 | 0.493 | 0.105 | - | 0.402 | 0.03 | 0.027 | - | 0.035 | 4.50E-25 | 1.15E-13 | 0.824 | 0.895 |
| *Sicily_EBA8561* | 0.141 | 0.601 | 0.165 | - | 0.233 | 0.028 | 0.026 | - | 0.035 | 1.65E-09 | 3.14E-09 | 0.141 | 0.191 |
| *Sicily_MBA3124* | 0.520 | 0.79 | 0.077 | - | 0.133 | 0.028 | 0.026 | - | 0.033 | 0.003 | 0.002 | 0.520 | 0.572 |
| *Sicily_MBA3125* | 0.330 | 0.771 | 0.049 | 0.180 | - | 0.040 | 0.021 | 0.036 | - | 7.20E-05 | 0.330 | 0.032 | 0.433 |
| *Sicily_MBA4109* | 0.243 | 0.778 | 0.074 | 0.149 | - | 0.044 | 0.022 | 0.039 | - | 0.003 | 0.243 | 0.037 | 0.324 |
| *Sicily_LBA* | 0.419 | 0.802 | 0.053 | - | 0.145 | 0.018 | 0.016 | - | 0.022 | 1.07E-08 | 2.47E-04 | 0.419 | 0.317 |
| *Sicilian (modern)* | 7.7E-06 | 0.458 | 0.046 | 0.354 | 0.141 | 0.022 | 0.014 | 0.028 | 0.023 | 9.57E-177 | 1.73E-11 | 2.07E-59 | 2.10E-09 |

* Note: For the analysis of the present-day Sardinians, we not only analyzed the 4 individuals with shotgun data (*Sardinian.DG*) but also 27 individuals with genotyping data on the Human Origins SNP array where we only have data at about half the positions but many more samples. For both the *Sardinian.DG* and the *Sardinian* analyses, the models only fit with *Iran_Ganj_Dareh_Neolithic* included as a source although with the larger number of samples that is available for the genotyping data we also have resolution to detect additional Steppe ancestry.

In **Supplementary Table 10** (and **Fig. 4**) we present these analyses on a by-individual basis. In what follows we selectively highlight some notable features of these analyses where they reveal patterns that are not clearly evidence in the grouped analysis.

For *Sardinia_Nuragic_BA*, individual I10554 produced the most parsimonious fitting model of 76.5 ± 2.7% *Anatolia_Neolithic*, 14.9 ± 2.7% *WHG*, and 8.5 ± 3.3% *Yamnaya_Samara* (p=0.16). The 2-way model with *Anatolia_Neolithic* and *WHG*, seen in the other Nuragic individuals, is weakly rejected at p=0.016, suggesting the possibility of a small proportion of Steppe ancestry in this individual. We view this inference with caution, however, as this individual lumps with the other *Sardinia_Nuragic_BA* individuals in *qpWave*, so any analysis of this individual alone is compromised by concerns about multiple hypothesis testing.

For *Sardinia_LateAntiquity*, one individual (I12222) failed all models we tested. However, when we removed *WHG*, we obtained a p=0.020 for a model of mixture of *Anatolia_Neolithic* and *Iran_Ganj_Dareh_Neolithic* (**Supplementary Table 10**). While below 0.050, this model is only a moderately poor fit, and makes sense in light of the fact that in PCA it clusters close to the other two individuals and close to *Anatolia_EBA* (**Fig. 2**).

For the *Sicily_EBA* cluster, 4 out of 5 individuals produce valid 2-way models with *Anatolia_Neolithic* and *WHG*, while I11442 fits a single 3-way model (p=0.391) with 69.0 ± 4.1% *Anatolia_Neolithic*, 11.6 ± 2.2% *WHG*, and 19.3 ± 3.8% *Iran_Ganj_Dareh_Neolithic* (*qpAdm* rejects the model of only *Yamnaya_Samara*-related ancestry for this individual (p=0.010)). The finding of Iranian-related ancestry in I11442 suggests that the Iranian-related ancestry that we detected in the Middle Bronze Age individuals may already have been present in the Early Bronze Age. However, we view this signal with caution as the p=0.010 rejection of *Yamnaya_Samara* model is only weak, and the *qpWave* clustering lumped this individual with the main *Sicily_EBA* group. Thus, the results of this by-individual analysis may be an artifact of multiple hypothesis testing.

The fact that that none of the models for *Sicily_EBA* individuals analyzed by themselves requires *Yamnaya_Samara* ancestry in order to fit, despite the fact that the analysis of the pool of all samples in **Supplementary Table 9** requires *Yamnaya_Samara*,is also notable. It highlights the statistical power that comes from pooling samples.

**Supplementary Table 10:** Admixture proportions for the most parsimonious *qpAdm* models of the all ancient individuals from this study fit as derived from groups related to four sources - *Anatolia_Neolithic* (1), *WHG* (2), *Iran_Ganj_Dareh_Neolithic* (3), and *Yamnaya_Samara* (4). In the final column, we show the p-value of the most parsimonious model (shaded in green) when we add *Morocco_LN* to the "Right" outgroup set. "Right" outgroup set: *Mbuti.DG, Ust_Ishim, CHG, EHG, ElMiron, Vestonice16, MA1, Israel_Natufian, Jordan_PPNB*. These data are used to produce **Fig. 4a**.

| Test | P-value | Admixture Sources and Proportions | | | | Standard Error | | | | P-values for other models | | | P-value of the most parsimonious model after adding Morocco_LN to the "Right" |
|---|---|---|---|---|---|---|---|---|---|---|---|---|---|
| | | 1 | 2 | 3 | 4 | SE1 | SE2 | SE3 | SE4 | 1+2 | 1+2+3 | 1+2+4 | |
| *Mallorca_EBA* | 0.665 | 0.454 | 0.177 | - | 0.369 | 0.036 | 0.033 | - | 0.042 | 0.000 | 0.000 | 0.665 | 0.758 |
| *Formentera_MBA* | 0.550 | 0.597 | 0.140 | - | 0.263 | 0.044 | 0.038 | - | 0.051 | 0.000 | 0.002 | 0.550 | 0.629 |
| *Menorca_LBA* | 0.299 | 0.594 | 0.174 | - | 0.231 | 0.028 | 0.027 | - | 0.036 | 0.000 | 0.000 | 0.299 | 0.370 |
| *Sardinia_Nuragic_BA3642* | 0.202 | 0.869 | 0.131 | | | 0.036 | 0.036 | | | 0.202 | 0.148 | 0.163 | 0.267 |
| *Sardinia_Nuragic_BA3741* | 0.173 | 0.842 | 0.158 | | | 0.022 | 0.022 | | | 0.173 | 0.470 | 0.114 | 0.228 |
| *Sardinia_Nuragic_BA3743* | 0.362 | 0.848 | 0.152 | | | 0.022 | 0.022 | | | 0.362 | 0.265 | 0.349 | 0.448 |
| *Sardinia_Nuragic_BA10364* | 0.214 | 0.795 | 0.205 | | | 0.022 | 0.022 | | | 0.214 | 0.357 | 0.362 | 0.262 |
| *Sardinia_Nuragic_BA10552* | 0.238 | 0.825 | 0.175 | | | 0.022 | 0.022 | | | 0.238 | 0.295 | 0.199 | 0.041 |
| *Sardinia_Nuragic_BA10553* | 0.208 | 0.804 | 0.196 | | | 0.022 | 0.022 | | | 0.208 | 0.230 | 0.184 | 0.217 |
| *Sardinia_Nuragic_BA10554* | 0.158 | 0.765 | 0.149 | | 0.085 | 0.027 | 0.027 | | 0.033 | 0.016 | 0.000 | 0.158 | 0.010 |
| *Sardinia_Nuragic_BA10365* | 0.064 | 0.854 | 0.146 | | | 0.022 | 0.022 | | | 0.064 | 0.389 | 0.436 | 0.099 |
| *Sardinia_IA10366* | 0.168 | 0.759 | 0.122 | 0.119 | - | 0.040 | 0.022 | 0.037 | - | 0.007 | 0.168 | 0.037 | 0.236 |
| *Sardinia_LateAntiquity12220* | 0.196 | 0.667 | 0.053 | 0.281 | - | 0.058 | 0.029 | 0.051 | - | 3.00E-06 | 0.196 | 4.30E-05 | 0.215 |
| *Sardinia_LateAntiquity12221* | 0.067 | 0.667 | - | - | 0.333 | 0.055 | - | - | 0.055 | 3.71E-07 | 0.001 | 0.243 | 0.097 |
| *Sardinia_LateAntiquity12222* | 0.020 | 0.674 | - | 0.326 | - | 0.083 | - | 0.083 | - | 1.00E-05 | 0.021 | 1.46E-04 | 0.033 |
| *Sardinia_LateAntiquity12223* | 0.328 | 0.580 | 0.057 | 0.362 | - | 0.119 | 0.050 | 0.101 | - | 0.004 | 0.328 | 0.049 | 0.382 |
| *Sicily_MN4062* | 0.386 | 0.900 | 0.100 | - | - | 0.022 | 0.022 | | | 0.386 | 0.288 | 0.379 | 0.412 |
| *Sicily_MN4063* | 0.243 | 0.914 | 0.086 | - | - | 0.021 | 0.021 | | | 0.243 | 0.230 | 0.257 | 0.321 |
| *Sicily_MN4065* | 0.146 | 0.825 | 0.175 | - | - | 0.023 | 0.023 | | | 0.146 | 0.097 | 0.099 | 0.185 |
| *Sicily_MN10376* | 0.100 | 0.928 | 0.072 | - | - | 0.022 | 0.022 | | | 0.100 | 0.063 | 0.066 | 0.151 |
| *Sicily_Bell_Beaker4936* | 0.474 | 0.831 | 0.169 | - | - | 0.079 | 0.079 | - | - | 0.474 | 0.374 | 0.265 | 0.395 |
| *Sicily_EBA3122* | 0.230 | 0.883 | 0.117 | - | - | 0.022 | 0.022 | | | 0.230 | 0.342 | 0.838 | 0.268 |
| *Sicily_EBA3123* | 0.623 | 0.752 | 0.107 | - | 0.141 | 0.029 | 0.025 | | 0.034 | 0.003 | 0.015 | 0.623 | 0.156 |
| *Sicily_EBA7796* | 0.389 | 0.923 | 0.077 | - | - | 0.031 | 0.031 | | | 0.389 | 0.468 | 0.685 | 0.431 |
| *Sicily_EBA7800* | 0.911 | 0.868 | 0.132 | - | | 0.055 | 0.055 | | | 0.911 | 0.978 | 0.978 | 0.871 |
| *Sicily_EBA7807* | 0.678 | 0.926 | 0.074 | - | - | 0.030 | 0.030 | | | 0.678 | 0.624 | 0.775 | 0.622 |
| *Sicily_EBA11442* | 0.391 | 0.690 | 0.116 | 0.193 | - | 0.041 | 0.022 | 0.038 | | 0.000 | 0.391 | 0.010 | 0.292 |
| *Sicily_EBA11443* | 0.824 | 0.493 | 0.105 | - | 0.402 | 0.030 | 0.027 | - | 0.035 | 0.000 | 0.000 | 0.824 | 0.895 |
| *Sicily_EBA8561* | 0.141 | 0.601 | 0.165 | - | 0.233 | 0.028 | 0.026 | - | 0.035 | 0.000 | 0.000 | 0.141 | 0.191 |
| *Sicily_MBA3124* | 0.520 | 0.79 | 0.077 | - | 0.133 | 0.028 | 0.026 | - | 0.033 | 0.003 | 0.002 | 0.520 | 0.572 |
| *Sicily_MBA3125* | 0.330 | 0.771 | 0.049 | 0.180 | - | 0.040 | 0.021 | 0.036 | - | 0.000 | 0.330 | 0.032 | 0.433 |
| *Sicily_MBA4109* | 0.243 | 0.778 | 0.074 | 0.149 | - | 0.044 | 0.022 | 0.039 | - | 0.003 | 0.243 | 0.037 | 0.324 |
| *Sicily_LBA3876* | 0.687 | 0.780 | 0.033 | - | 0.186 | 0.028 | 0.027 | - | 0.035 | 0.000 | 0.000 | 0.687 | 0.553 |
| *Sicily_LBA3878 (fit 1)* | 0.594 | 0.817 | 0.062 | 0.122 | - | 0.026 | 0.026 | 0.032 | - | 0.019 | 0.805 | 0.594 | 0.400 |
| *Sicily_LBA3878 (fit 2)* | 0.805 | 0.745 | 0.119 | - | 0.135 | 0.040 | 0.021 | - | 0.036 | 0.019 | 0.805 | 0.594 | 0.328 |
| *Sicily_LBA10372* | 0.045 | 0.797 | 0.076 | - | 0.127 | 0.028 | 0.028 | - | 0.034 | 0.000 | 0.004 | 0.045 | 0.052 |
| *Sicily_LBA10373 (fit 1)* | 0.201 | 0.762 | 0.106 | 0.132 | - | 0.042 | 0.021 | 0.039 | - | 0.010 | 0.201 | 0.485 | 0.229 |
| *Sicily_LBA10373 (fit 2)* | 0.485 | 0.820 | 0.047 | - | 0.133 | 0.028 | 0.026 | - | 0.034 | 0.010 | 0.201 | 0.485 | 0.497 |
| *Sicily_LBA10371* | 0.285 | 1.040 | -0.040 | - | - | 0.053 | 0.053 | - | - | 0.285 | 0.202 | 0.207 | 0.343 |

* For *Sardinia_LateAntiquity12221* and *Sardinia_LateAntiquity12222*, the only fitting three-way model was *Anatolia_Neolithic+WHG+Yamnaya_Samara*, but the estimates of *WHG* were negative so at -11.9 ± 5.4% so we show the model without WHG ancestry which also fit (at p=0.067).

##### 3.2 - Results of Proximal Modelling

The objective of proximal *qpAdm* modelling is to identify possible sources of admixture that are more closely related in geography and time to a *Test* population. As a base "Right" set for the proximal modelling we used the same Right population that were used for the distal modeling, but we now also adding in the four sources of the distal modeling.

*Mbuti.DG, Ust_Ishim, CHG, EHG, ElMiron, Vestonice16, MA1, Israel_Natufian, Jordan_PPNB, Anatolia_Neolithic, WHG, Iran_Ganj_Dareh_Chalcolithic, Yamnaya_Samara*

We tested different relevant sets of sources for each region and followed a similar approach as in [33]. We first identified all valid models with the fixed base "Right" set, and then we re-evaluated these by adding the remaining unused sources to the "Right" set of populations ("model competition"). The results presented are after "model competition". We considered models as valid if their fit was p>0.05.

###### Balearic Bronze Age Proximal Modeling

To investigate the origins and ancestry of the three Balearic Bronze Age individuals we tested the following populations as sources, examining all possible combinations of 1-, 2-, and 3-way models, and highlighting the parsimonious ones:

*Bell_Beaker_Iberia, Bell_Beaker_Iberia_highsteppe, Iberia_Chalcolithic, France_Bell_Beaker, France_Bell_Beaker_lowsteppe, Sardinia_Nuragic_BA*

*Mallorca_EBA*: We found a single valid 1-way model with *Bell_Beaker_Iberia_highsteppe*, a group of outliers from Iberia buried in a Bell Beaker mortuary context who unlike most individuals from this context in that region had high proportions of Steppe ancestry (p=0.442)[41].

*Formentera_MBA*: We added *Mallorca_EBA* as a possible source, and found that the 1-way models with *Mallorca_EBA* (p=0.774) and *Bell_Beaker_Iberia_highsteppe* (0.533) were fits.

*Menorca_LBA*: We added both *Mallorca_EBA* and *Formentera_MBA* as a possible sources. The only parsimonious 1-way model was *Formentera_MBA* (p=0.531).

*Ibiza_Phoenician*: We investigated if the published Phoenician individual from Ibiza[32] was consistent with inheriting some ancestry from previous Balearic Islands populations so we used the same proximal sources as for *Menorca_LBA* but then added: *Menorca_LBA*, *Mycenaean*, *Sardinia_IA*, *Sicily_MBA4109*, *Morocco_LN*, and *Jordan_EBA*. Only models with two sources of admixture produced valid results, and all of them required *Morocco_LN* as one of those sources (**Supplementary Table 11**). Even though we used model competition to try to reduce the number of working models by adding the unused sources to the "Right" (including the Bronze Age Balearic individuals), none of the initially working models failed. Considering that in *qpWave* this individual did not form a clade with the other Balearic it is possible that these models represent an unsampled population. These results clearly demonstrate a link to North African ancestry in the Phoenician settlement of the Balearic Islands.

**Supplementary Table 11:** Most parsimonious *qpAdm* models for *Ibiza_Phoenician*.

| Admixture Sources | | P-value | Mixture Proportion | | Standard Error | |
|---|---|---|---|---|---|---|
| A | B | | Anc_A | Anc_B | SE_A | SE_B |
| *Bell_Beaker_Iberia_highsteppe* | *Morocco_LN* | 0.518 | 0.146 | 0.854 | 0.036 | 0.036 |
| *Bell_Beaker_Iberia* | *Morocco_LN* | 0.130 | 0.090 | 0.910 | 0.034 | 0.034 |
| *Formentera_MBA* | *Morocco_LN* | 0.468 | 0.434 | 0.566 | 0.116 | 0.116 |
| *France_Bell_Beaker* | *Morocco_LN* | 0.869 | 0.171 | 0.829 | 0.035 | 0.035 |
| *Mallorca_EBA* | *Morocco_LN* | 0.732 | 0.343 | 0.657 | 0.076 | 0.076 |
| *Menorca_LBA* | *Morocco_LN* | 0.531 | 0.167 | 0.833 | 0.045 | 0.045 |
| *Mycenaean* | *Morocco_LN* | 0.099 | 0.251 | 0.749 | 0.116 | 0.116 |

**Sardinian Bronze Age Proximal Modeling**

*Sardinia_Nuragic_BA*: Although the individuals within this group date to a broad temporal range from 2134-1947 calBCE (I3743) to 1116-824 calBCE (I10364), they all form a clade with similar genetic composition. To better understand the ancestry of this grouping, we tested as sources ancient groups chronologically close to the oldest Nuragic Bronze Age individuals (~2000 BCE) that would allow for the identification of both Steppe and Iranian-related ancestries. We also included as a possible source *Sicily_MN*, which shares a similar ancestry profile in ADMIXTURE (**Fig. 2**):

*Bell_Beaker_Iberia, Bell_Beaker_Iberia_highsteppe, Czech_Bell_Beaker, Iberia_Chalcolithic, France_Bell_Beaker, France_Bell_Beaker_lowsteppe, Sicily_MN, Mallorca_EBA, Anatolia_EBA*

No model fit $p>0.05$ (or even $p>0.01$), suggesting that the early European farmer-related ancestry in Sardinia was distinct from that in the other places in Europe for which we have data.

*Sardinia_Nuragic_BA10365*: For the Middle Bronze Age outlier I10365 (1643-1263 calBCE) we added *Sicily_EBA*, *Sardinia_Nuragic_BA*, *Minoan_Lassithi* and *Mycenaean* as additional possible sources to the ones also tested for the larger *Sardinia_Nuragic_BA* grouping. The results, shown in **Supplementary Table 12**, reveal four parsimonious 2-way admixture models that always require *Sardinia_Nuragic_BA* (suggesting some degree of local continuity) and either a source of Iranian-related ancestry (*Anatolia_EBA* or *Mycenaean*), or Steppe-related ancestry (*France_Bell_Beaker* or *Czech_Bell_Beaker*). The inference of eastern ancestry in this individual is different from the finding in the distal modeling where a model of just *Anatolia_Neolithic* and *WHG* passes ($p=0.064$), although this is in no way a contradiction because the proximal modeling potentially has more statistical power. As documented above, modern Sardinians definitively do have eastern admixture, which may explain why present-day Sardinians are consistent in our *qpWave* analysis with forming a clade *with Sardinia_Nuragic_BA10365* but not with the main *Sardinia_Nuragic_BA* cluster.

**Supplementary Table 12:** Most parsimonious *qpAdm* models for *Sardinia_Nuragic_BA10365*.

| Admixture Sources | | P-value | Mixture Proportion | | Standard Error | |
|---|---|---|---|---|---|---|
| A | B | | Anc_A | Anc_B | SE_A | SE_B |
| *Sardinia_Nuragic_BA* | *Anatolia_EBA* | 0.323 | 0.779 | 0.221 | 0.043 | 0.043 |
| *Sardinia_Nuragic_BA* | *Mycenaean* | 0.517 | 0.604 | 0.396 | 0.065 | 0.065 |
| *Sardinia_Nuragic_BA* | *Czech_Bell_Beaker* | 0.565 | 0.802 | 0.198 | 0.035 | 0.035 |
| *Sardinia_Nuragic_BA* | *France_Bell_Beaker* | 0.298 | 0.847 | 0.153 | 0.031 | 0.031 |

*Sardinia_IA*:

For the Iron Age individual we further included *Jordan_EBA* as a potential source. We found one valid model for *Iberia_Chalcolithic* (11.9 ± 3.2%) and *Mycenaean* (88.1 ± 3.2%) (p=0.067). Notably, this model works even with inclusion of *Sardinia_Nuragic_BA* in the outgroup set, and thus with our analysis we cannot rule out the possibility of very little contribution of Nuragic Sardinians to the Iron Age Sardinian. The addition of *Morocco_LN* to the "Right" does not break the model (p=0.097), so there is no evidence for North African ancestry in this individual either.

*Sardinia_LateAntiquity*:

For the main group of 3 *Sardinia_LateAntiquity* individuals we explored the following as possible sources:

*Croatia_Early_IA, Hungary_Prescythian_IA.SG, Bulgaria_IA, Egypt_Hellenistic, Anatolia_Hellenistic.SG, Morocco_LN, Hungary_Scythian.SG, Jordan_EBA, Russia_Sarmatian.SG, Sardinia_Nuragic_BA, Sardinia_Nuragic_BA10365, Russia_Scythian_IA, Sardinia_IA, Mycenaean, Ibiza_Phoenician*

We found that we required only 1-way models to produce a valid p>0.05, obtaining fits for both *Mycenaean* (p=0.241) and *Ibiza_Phoenician* (p=0.145). Notably, both these fits work with *Sardnia_Nuragic_BA* among the outgroups, showing that there is no evidence of these Late Antiquity individuals inheriting any ancestry from Nurgaric Bronze Age Sardinians.

*Sardinia_LateAntiquity12221*:

We used the same sources for *Sardinia_LateAntiquity12221* and observed many working 2-way models, possibly reflecting the limit power of this analysis with a single individual of modest coverage (78437 autosomal SNPs) (**Supplementary Table 13**). The lack of a historically plausible pair of source populations is again likely to reflect lack of available data from this region. We ote that many of these models do not include Sardinian source populations, and all these models include Nuragic Bronze Age Sardinians in the outgroups. Thus, *Sardinia_LateAntiquity12221,* like the main *Sardinia_LateAntiquity* genetic cluster, has no evidence of ancestry from early Sardinians.

**Supplementary Table 13:** Most parsimonious *qpAdm* models for *Sardinia_LateAntiquity*.

| Admixture Sources | | P-value | Mixture Proportion | | Standard Error | |
|---|---|---|---|---|---|---|
| A | B | | Anc_A | Anc_B | SE_A | SE_B |
| *Anatolia_Hellenistic.SG* | *Sardinia_IA* | 0.836 | 0.597 | 0.403 | 0.122 | 0.122 |
| *Anatolia_Hellenistic.SG* | *Sardinia_Nuragic_BA10365* | 0.437 | 0.717 | 0.283 | 0.141 | 0.141 |
| *Anatolia_Hellenistic.SG* | *Sardinia_Nuragic_BA* | 0.926 | 0.696 | 0.304 | 0.067 | 0.067 |
| *Bulgaria_IA* | *Anatolia_Hellenistic.SG* | 0.134 | 0.174 | 0.826 | 0.174 | 0.174 |
| *Bulgaria_IA* | *Hungary_Scythian.SG* | 0.111 | 0.376 | 0.624 | 0.126 | 0.126 |
| *Croatia_Early_IA* | *Egypt_Hellenistic* | 0.480 | 0.466 | 0.534 | 0.168 | 0.168 |
| *Croatia_Early_IA* | *Morocco_LN* | 0.843 | 0.412 | 0.588 | 0.102 | 0.102 |
| *Egypt_Hellenistic* | *Hungary_Scythian.SG* | 0.299 | 0.66 | 0.340 | 0.125 | 0.125 |
| *Egypt_Hellenistic* | *Sardinia_Nuragic_BA10365* | 0.212 | 0.710 | 0.290 | 0.124 | 0.124 |
| *Egypt_Hellenistic* | *Sardinia_Nuragic_BA* | 0.061 | 0.784 | 0.216 | 0.088 | 0.088 |
| *Hungary_Prescythian_IA.SG* | *Bulgaria_IA* | 0.176 | 0.487 | 0.513 | 0.089 | 0.089 |
| *Hungary_Prescythian_IA.SG* | *Morocco_LN* | 0.510 | 0.296 | 0.704 | 0.065 | 0.065 |
| *Hungary_Prescythian_IA.SG* | *Sardinia_IA* | 0.595 | 0.368 | 0.632 | 0.099 | 0.099 |
| *Hungary_Prescythian_IA.SG* | *Sardinia_Nuragic_BA10365* | 0.058 | 0.436 | 0.564 | 0.098 | 0.098 |
| *Hungary_Prescythian_IA.SG* | *Sardinia_Nuragic_BA* | 0.298 | 0.541 | 0.459 | 0.067 | 0.067 |
| *Hungary_Scythian.SG* | *Jordan_EBA* | 0.770 | 0.615 | 0.385 | 0.074 | 0.074 |
| *Hungary_Scythian.SG* | *Sardinia_IA* | 0.258 | 0.423 | 0.577 | 0.169 | 0.169 |
| *Morocco_LN* | *Hungary_Scythian.SG* | 0.730 | 0.630 | 0.370 | 0.080 | 0.080 |
| *Morocco_LN* | *Russia_Sarmatian.SG* | 0.204 | 0.789 | 0.211 | 0.054 | 0.054 |
| *Russia_Sarmatian.SG* | *Sardinia_IA* | 0.615 | 0.294 | 0.706 | 0.072 | 0.072 |
| *Russia_Sarmatian.SG* | *Sardinia_Nuragic_BA10365* | 0.083 | 0.350 | 0.650 | 0.065 | 0.065 |
| *Russia_Scythian_IA* | *Morocco_LN* | 0.421 | 0.195 | 0.805 | 0.042 | 0.042 |
| *Russia_Scythian_IA* | *Sardinia_IA* | 0.602 | 0.255 | 0.745 | 0.064 | 0.064 |

*Sardinian*: For proximal modeling of modern Sardinians we used the same sources as for *Sardinia_LateAntiquity* and *Sardinia_LateAntiquity12221* but here included them as potential sources as well. After model competition only one model remained significant (p=0.433), even when a lower threshold of p>0.01 is applied, with 13.6 ± 3.4% *Sardinia_Nuragic_BA* and 86.4 ± 3.4% *Sardinia_LateAntiquity12221.*

**Sicilian Bronze Age Proximal Modeling**

We modeled Bronze Age Sicilians as having a local source of ancestry represented by *Sicily_MN*, and then added combinations of Bell Beaker culture associated populations from the west (as we find Iberia-specific Y chromosome haplogroup R1b1a1a2a1a2a1 (Z195) in Early Bronze Age Sicily), namely Iberia and France (*Bell_Beaker_Iberia_highsteppe, France_Bell_Beaker).* We also use *Minoan_Lassithi* as a proxy for Iranian-related ancestry.

*Early Bronze Age*: We find that *Sicily_EBA* can be fit as *Sicily_MN* and *Bell_Beaker_Iberia_highsteppe* (15.7 ± 3.5%), in agreement with the Y chromosome evidence of an Iberian affinity. Our modeling of the three outliers identifies *France_Bell_Beaker* as the only parsimonious fitting second source (**Supplementary Table 14**), albeit with different proportions of 29.7 ± 3.4% in *Sicily_EBA3123*, 44.7 ± 3.2% in *Sicily_EBA8561*, and , 74.2 ± 3.8 in *Sicily_EBA11443*.

**Supplementary Table 14:** Most parsimonious *qpAdm* models for Early Bronze Age Sicilians.

| *Test* | Admixture Sources | | P-value | Mixture Proportion | | Standard Error | |
|---|---|---|---|---|---|---|---|
| | A | B | | Anc_A | Anc_B | SE_A | SE_B |
| *Sicily_EBA* | *Sicily_MN* | *Bell_Beaker_Iberia_highsteppe* | 0.120 | 0.843 | 0.157 | 0.035 | 0.035 |
| *Sicily_EBA3123* | *Sicily_MN* | *France_Bell_Beaker* | 0.377 | 0.703 | 0.297 | 0.034 | 0.034 |
| *Sicily_EBA11443* | *Sicily_MN* | *France_Bell_Beaker* | 0.297 | 0.258 | 0.742 | 0.038 | 0.038 |
| *Sicily_EBA8561* | *Sicily_MN* | *France_Bell_Beaker* | 0.624 | 0.553 | 0.447 | 0.032 | 0.032 |

*Middle and Late Bronze Age*: If we do not include *Sicily_EBA* as a potential proximal source, two of the three Sicilian individuals dated to the Middle Bronze Age (I3125 and I4109) produce models that require both a source of Steppe ancestry and another of Iranian-related ancestry (**Supplementary Table 15**), consistent with the distal modeling results.

**Supplementary Table 15:** *qpAdm* models for Middle and Late Bronze Age Sicilians.

| *Test* | Admixture Sources | | | P-value | Mixture Proportion | | | Standard Error | | |
|---|---|---|---|---|---|---|---|---|---|---|
| | A | B | C | | Anc_A | Anc_B | Anc_C | SE_A | SE_B | SE_C |
| *Sicily_MBA3124* | *Sicily_MN* | *France_Bell_Beaker* | | 0.069 | 0.733 | 0.267 | - | 0.032 | 0.032 | - |
| *Sicily_MBA3125* | *Sicily_MN* | *Minoan_Lassithi* | *Bell_Beaker_Iberia_highsteppe* | 0.144 | 0.122 | 0.646 | 0.232 | 0.115 | 0.100 | 0.046 |
| | *Sicily_MN* | *Minoan_Lassithi* | *France_Bell_Beaker* | 0.092 | 0.318 | 0.529 | 0.152 | 0.100 | 0.101 | 0.031 |
| *Sicily_MBA4109* | *Sicily_MN* | *Minoan_Lassithi* | *Bell_Beaker_Iberia_highsteppe* | 0.294 | 0.352 | 0.549 | 0.100 | 0.128 | 0.111 | 0.049 |
| | *Sicily_MN* | *Minoan_Lassithi* | *France_Bell_Beaker* | 0.268 | 0.440 | 0.496 | 0.064 | 0.112 | 0.111 | 0.032 |
| *Sicily_LBA* | *Sicily_MN* | *Minoan_Lassithi* | *Bell_Beaker_Iberia_highsteppe* | 0.101 | 0.324 | 0.431 | 0.244 | 0.072 | 0.061 | 0.031 |
| | *Sicily_MN* | *Minoan_Lassithi* | *France_Bell_Beaker* | 0.197 | 0.530 | 0.303 | 0.167 | 0.062 | 0.062 | 0.021 |

When we add Early Bronze Age individuals as potential sources in the proximal modeling for the Middle Bronze Age Sicilians, *Sicily_EBA3123* fits as a source (without mixture) for *Sicily_MBA3124*, but we obtain no other fitting models for other Middle Bronze Age individuals.

When we add Early and Middle Bronze Age individuals as potential sources in the proximal modeling for the Late Bronze Age, we find that *Sicily_EBA3123, Sicily_MBA3124,* and *Sicily_MBA3125* all fit as a single source (without mixture) for *Sicily_LBA,* and hence we do not show a table of fits.
